# Natural combinatorial genetics and prolific polyamine production enable siderophore diversification in Serratia plymuthica

**DOI:** 10.1101/2020.07.03.184812

**Authors:** Sara Cleto, Kristina Haslinger, Kristala L.J. Prather, Timothy K. Lu

**Affiliations:** Department of Electrical Engineering and Computer Science, Massachusetts Institute of Technology; Department of Biological Engineering, Massachusetts Institute of Technology; Synthetic Biology Center, Massachusetts Institute of Technology; Department of Chemical Engineering, Massachusetts Institute of Technology; Department of Chemical and Pharmaceutical Biology, University of Groningen

**Keywords:** Natural products, siderophores, pathway elucidation, polyamines

## Abstract

Siderophores are small molecules with unmatched capacity to scavenge iron from proteins and the extracellular milieu, where it mostly occurs as insoluble Fe^3+^. Siderophores chelate Fe^3+^ for uptake into the cell, where it is reduced to soluble Fe^2+^. As iron is essential for bacterial survival, siderophores are key molecules in low soluble iron conditions. Bacteria have devised many strategies to synthesize proprietary siderophores to avoid siderophore piracy by competing organisms, e.g., by incorporating different polyamine backbones into siderophores, while maintaining the catechol moieties. We report that *Serratia plymuthica* V4 produces a variety of siderophores, which we term the *siderome*, and which are assembled by the concerted action of enzymes encoded in two independent gene clusters. Besides assembling serratiochelin with diaminopropane, *S. plymuthica* utilizes putrescine and the same set of enzymes to assemble photobactin, a siderophore described for *Photorhabdus luminescens*. The enzymes encoded by one of the gene clusters can independently assemble enterobactin. A third, independent operon is responsible for biosynthesis of the hydroxamate siderophore aerobactin, initially described in *Enterobacter aerogenes*. Mutant strains not synthesizing polyamine-siderophores significantly increased enterobactin production levels, though lack of enterobactin did not impact serratiochelin production. Knocking out SchF0, an enzyme involved in the assembly of enterobactin alone, significantly reduced bacterial fitness. This study illuminates the interplay between siderophore biosynthetic pathways and polyamine production superpathways, indicating routes of molecular diversification. Given its natural yields of diaminopropane (97.75 μmol/g DW) and putrescine (30.83 μmol/g DW), *S. plymuthica* can be exploited for the industrial production of these compounds.

**Significance Statement:** Siderophores are molecules crucial for bacterial survival in low iron environments. Bacteria have evolved the capacity to pirate siderophores made by other bacterial strains and to diversify the structure of their own siderophores, to prevent piracy. We found that *Serratia plymuthica* V4 produces five different siderophores using three gene clusters and a polyamine production superpathway. The most well studied siderophore, enterobactin, rather than the strain’s proprietary and by far most abundant siderophore, serratiochelin, displayed a crucial role in the fitness of *S. plymuthica*. Our results also indicate that this strain is a good candidate for engineering the large-scale production of diaminopropane (DAP), as without any optimization it produced the highest amounts of DAP reported for wild-type strains.

## Introduction

Iron, one of the most abundant elements on Earth (1), is crucial for the survival of all living organisms, including bacteria. It occurs in two forms: soluble (Fe^2+^) and insoluble (Fe^3+^). Soluble iron can be readily taken up by aerobic microorganisms (but not anaerobes), although it is uncommon at pH 7 (2–4). Bacteria and most life forms have evolved a diversity of ways that converge to the same goal: obtaining soluble iron (Fe^2+^) for survival. They have devised complex regulatory mechanisms responding to Fe^2+^ unavailability that induce the expression of a series of genes to produce small iron chelators, termed siderophores (5–8), secrete them, and take up their iron-bound forms. Bacteria have not only devised ways of biosynthesizing “proprietary” siderophore molecules, but have evolved transport mechanisms that allow them to utilize foreign siderophores, or xenosiderophores, as well (9, 10). This mechanism has led siderophores to be considered public goods, traded between bacteria and impacting their survival (11–15). As with any public good, some users benefit from it without having contributed to its production, which comes at a cost to the producer (15). Along these lines, some bacteria have evolved extraordinary ways to synthesize proprietary siderophores that require the expression of specialized TonB-dependent receptors (TBDRs). These receptors allow for efficient siderophore uptake by the producer: competitors lacking the receptor cannot take those siderophores up; thus, no piracy can occur (16, 17). One such innovative way is the incorporation of polyamines into the nascent siderophore, which has evolved in multiple species that naturally produce polyamines. Thus, diaminopropane (DAP) is incorporated into serratiochelin in *Serratia plymuthica* (18), norspermidine is incorporated into vibriobactin in *Vibrio cholerae* (19) and vulnibactin in *Vibrio vulnificus* (20), putrescine is incorporated into photobactin in *Photorhabdus luminescens* (21), and spermidine is incorporated into parabactin in *Paracoccus denitrificans* (22) and agrobactin in *Agrobacterium tumefaciens* (23).

Polyamines are small organic molecules with various numbers of carbons and amine moieties and a flexible structure (24, 25). They are synthesized by most bacteria and some eukaryotes (26) from L-lysine, L-methionine, L-aspartate, and L-arginine, with bacteria synthesizing a greater diversity of polyamines than eukaryotes. These moieties are incorporated into the nascent siderophore molecules by dedicated amide synthases, which contain stand-alone condensation domains structurally related to those found in non-ribosomal peptide synthetases (27, 28). Amide synthases have already been identified in several organisms that produce polyamine-containing siderophores, such as PhbG in *Photorhabdus* spp. (21), SchH in *Serratia* spp. (18), and especially VibH in *Vibrio* spp. (28). VibH has been crystalized and the condensing activity thoroughly studied (28). The amide synthase involved in the assembly of agrobactin in *Agrobacterium* spp. has yet to be identified though its biosynthetic cluster is known (29). The biosynthetic cluster for parabactin, thus also its amide synthase, has yet to be identified.

*S. plymuthica* stands out for its ability to produce the nonribosomal peptide antibiotic zeamine and the nonribosomal peptide siderophore serratiochelin (30, 31). Gene clusters evolutionarily obtained by *S. plymuthica* from a diversity of bacteria, such as *Dickeya zeae* (30, 31), *Escherichia coli*, and *Vibrio* spp. (18), are involved in the assembly of these molecules. In this work, we further explored and elucidated the diversity of siderophores produced by *S. plymuthica* and dissected the interplay of two catechol siderophore pathways with a superpathway for polyamine production, as well as their role in the diversification of catechol siderophores in this organism. In addition, to shed light on the relationship between the amide synthases and their preference for specific polyamines, we identified active site residues using bioinformatics tools. Furthermore, we dissected the diversity of putative TBDRs in the genome of *S. plymuthica*.

In this work, *S. plymuthica* was found to produce an extraordinary diversity of siderophores, which we termed the *siderome*. This diversity is generated by an interplay of three independent siderophore biosynthetic clusters and a prolific polyamine production superpathway, which is rare among Enterobacteriaceae. These siderophores were serratiochelin, enterobactin, photobactin, and aerobactin. To the best of our knowledge, this is the first published natural occurrence of serratiochelin, photobactin, enterobactin, and aerobactin in a single bacterial species. These findings suggest that the capacity of *S. plymuthica* to accrue biosynthetic clusters that evolved in other organisms is more extensive than so far described. Our results emphasize the utility of studying the evolution of natural product biosynthetic pathways and networks.

## Results

### Characterization of the siderophores produced by *S. plymuthica*

*S. plymuthica* produces serratiochelin (18), enterobactin (32, 33), photobactin (21), and aerobactin (34) (Figure 1). The presence and identity of the molecules was investigated using liquid chromatography coupled tandem mass spectrometry analyzed by XCMS, Thermo XCalibur^®^, PubChem, and Chemdraw, and by comparing the fragmentation pattern of each of the predicted or known patterns (depicted in Supplemental Figures 1 through 4). When the gene clusters involved in the assembly of serratiochelins were originally characterized (here p1 and p2, Figure 1b), it was found that the enzyme SchF0 encoded in gene cluster p1 and homologous to EntF *in E. coli*, was not involved in this process. Instead, the enzymes SchF1F2F3 encoded in gene cluster p2 and homologous to VibF in *V. cholerae*, were involved (18). Given that enterobactin is also synthesized by *S. plymuthica*, we hypothesized that although SchF0 is not involved in serratiochelin assembly, SchF0 might nonetheless be essential for enterobactin assembly. We thus screened wildtype and SchF0 mutants of *S. plymuthica* for the production of enterobactin (Figure 2d). We found that we observed that the disruption of *schF0* affected only the assembly of enterobactin, whereas the disruption of *schF3* abolished the production of all other catecholate siderophores (Figure 2). Accordingly, *schF0* is not a pseudogene as previously suggested (18) but a gene utilized specifically for the assembly of enterobactin and not for other catecholate siderophores in this organism.

**Figure 1.**
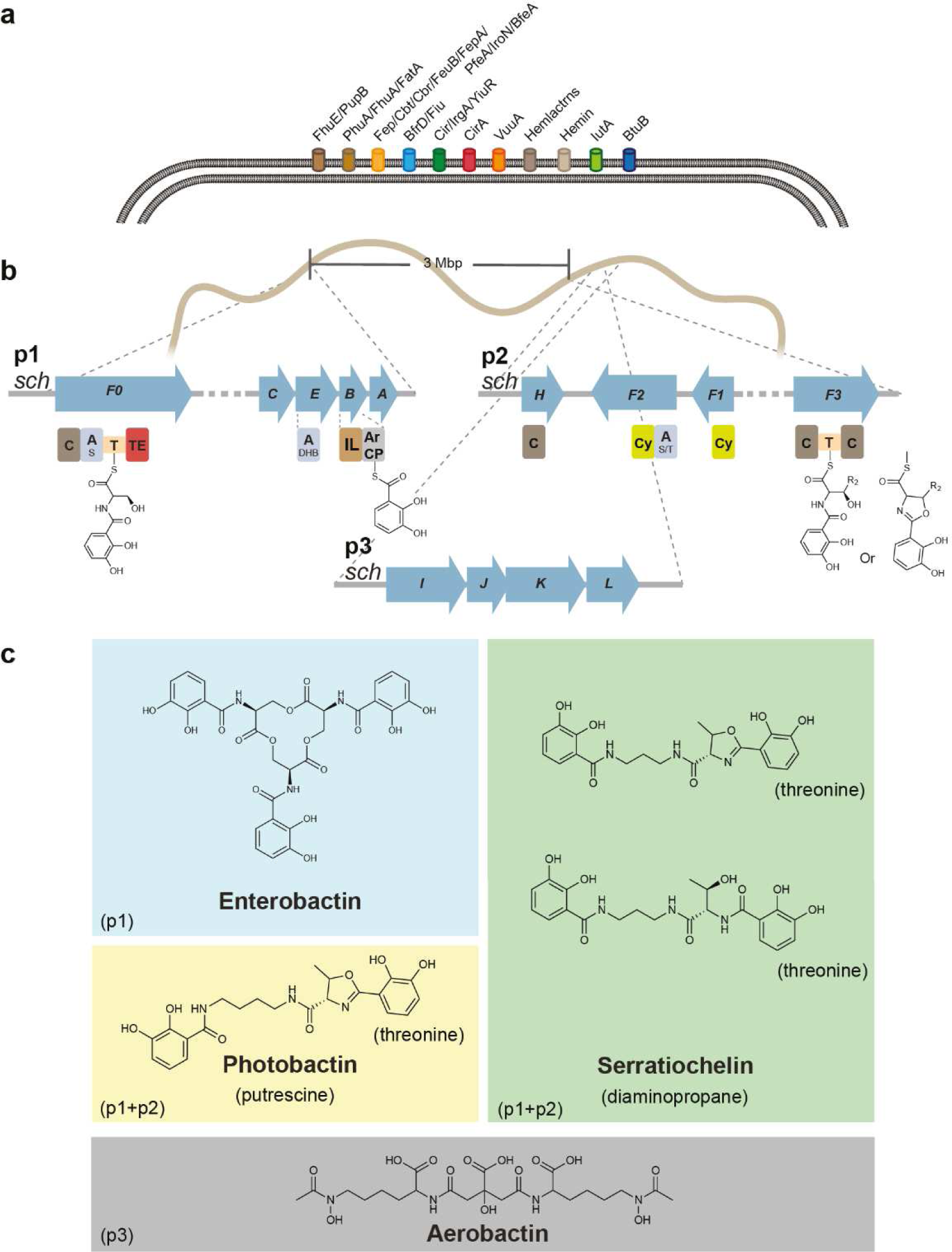
Schematic of the molecular players in the iron uptake mechanism of *S. plymuthica*. Putative TonB-dependent siderophore uptake receptors identified in the *S. plymuthica* genome (a); siderophore-encoding gene clusters (p1-p3) identified in the *S. plymuthica* genome and experimentally characterized in this study (domain annotations below blue arrow: C=condensation domain, A=adenylation domain (index indicates the substrate; 2,3-dihydroxybenzoate (DHB), serine (S), threonine (T)), T=thiolation domain, IL=isochorismate lyase ArCP=aryl carrier protein, TE=thioesterase domain, Cy=cyclisation domain, (b); siderophores detected in this study grouped by their underlying biosynthetic pathways (p1-p3)(c).

**Figure 2.**
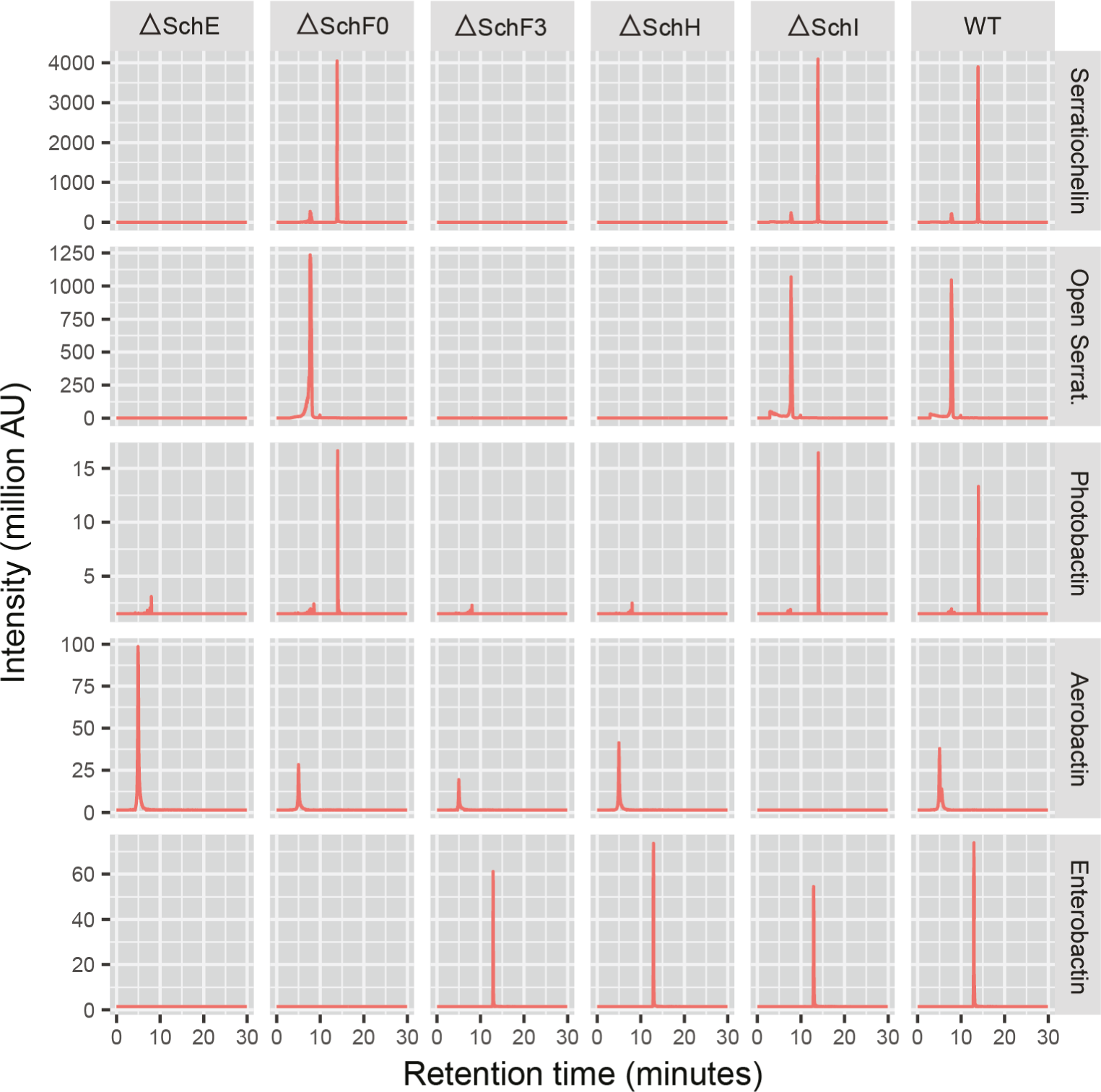
Extracted Ion Count in million activity units for serratiochelin (closed and open form), photobactin, aerobactin, and enterobactin in the *schE, schF0, schF3, schH, schI* mutants and the wild-type strain.

Next, we sought to characterize SchE encoded in gene cluster p2. This is a homologue of EntE, which adenylates the catechol precursor of enterobactin, 2,3-dihydroxybenzoate (DHB), and tethers it to holo-EntB (35). This is a crucial step in catecholate siderophore assembly (35). We screened an SchE mutant for the production of each of the aforementioned siderophores. As expected, solely aerobactin, which is a hydroxamate siderophore, was synthesized. This result agrees with our predictions and shows that no other EntE/SchE homologes are present in the genome of *S. plymuthica*. In an earlier study, it was found that the condensation of polyamines with DHB is catalyzed by SchH in *S. plymuthica* (encoded in cluster p2) (18). Therefore, we screened a SchH deletion mutant for the biosynthesis of serratiochelin and photobactin, which contain polyamine moieties (DAP and putrescine, respectively). We found that the SchH knockout strain did not synthesize these siderophores. Aerobactin was still produced, as was enterobactin. These siderophores do not have a polyamine moiety, and so the inability to synthesize the polyamine would not have affected the production of these siderophores. In fact, enterobactin was overproduced, in comparison with enterobactin production from the wild-type strain (p = 0.004).

Lastly, given that we detected aerobactin in our samples, we decided to query the genome of *S. plymuthica* for genes homologous to those involved in the biosynthesis of aerobactin in other organisms. This enabled us to locate a chromosomal operon homologous to *iucABCD*, which we termed *schIJKL* (p3), as not to be confused with *schABCD* (34) (Figure 3a). The four genes in this operon were highly similar to those in a *Yersinia* strain, with identities as high as 89%, as determined by pairwise analysis with the Basic Local Alignment Search Tool, BLAST2p (Supplemental Table 1). LucA (a homolog of SchI) catalyzes the intermediate step that converts L-lysine to aerobactin (34). To test whether this operon was indeed responsible for the production of the hydroxamate siderophore aerobactin, we built a SchI-defective mutant and tested it for the capacity to synthesize aerobactin. We found that the ΔSchI strain did not produce aerobactin (Figure 3d and e), whereas the wild-type strain and all other mutants were capable of synthesizing aerobactin (Figure 2). This confirms that the *schIJKL* operon (p3) is indeed responsible for the biosynthesis of aerobactin.

**Table 1.**
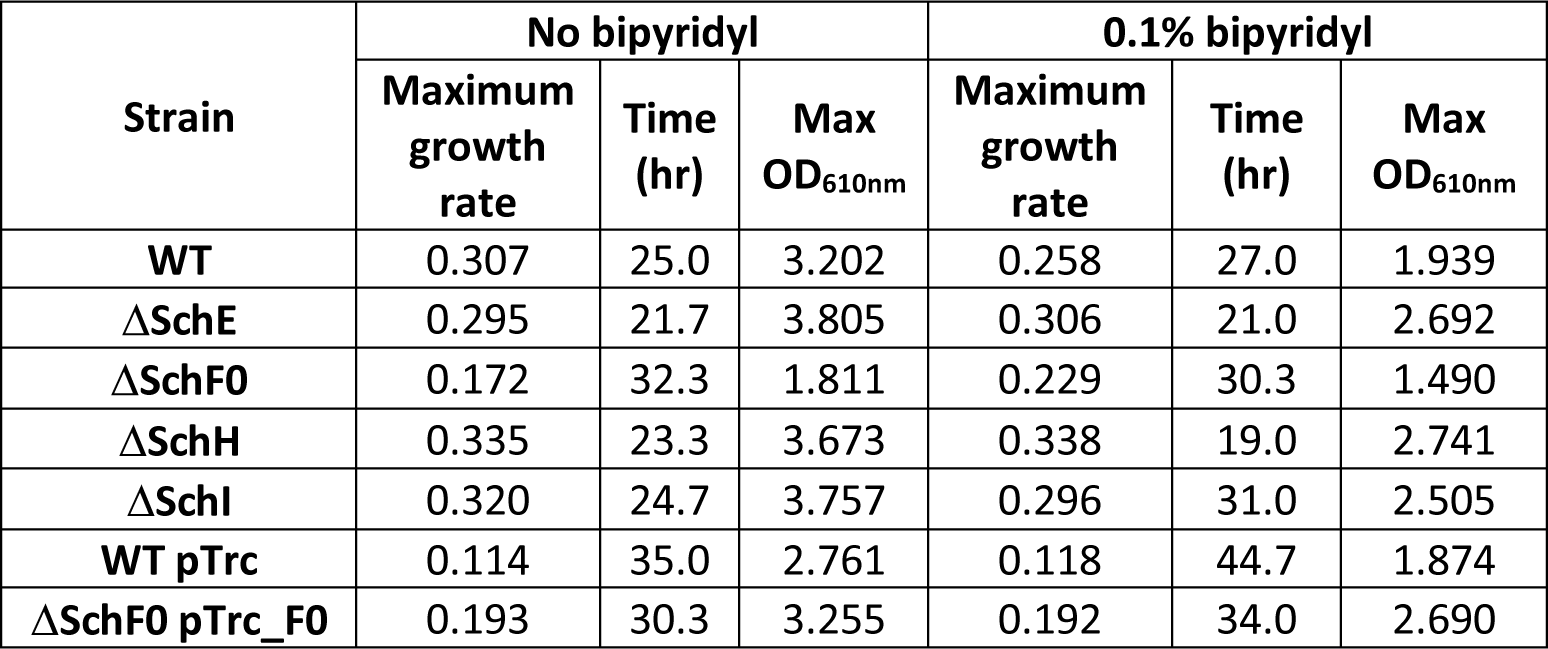
Maximum growth rates and OD_610nm_ of wild-type *S. plymuthica* and siderophore mutant strains grown in minimal medium, in the presence and absence of bipyridyl.

**Figure 3.**
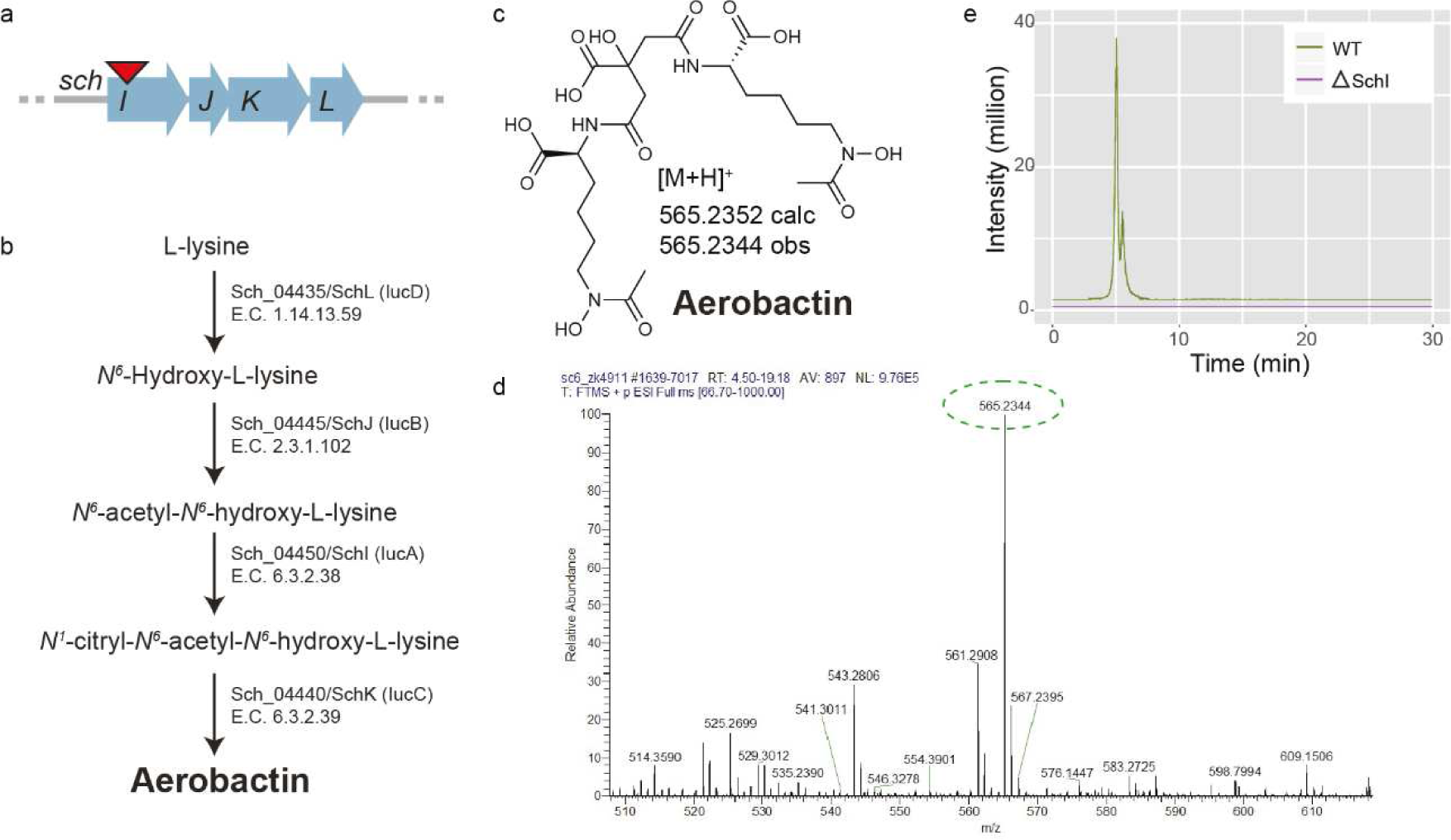
Characterization of aerobactin biosynthesis. Biosynthetic operon and respective locus of suicide vector introduction (a), enzymatic processes leading to aerobactin biosynthesis (b), calculated and observed mass of aerobactin (c), ESI-MS of wild-type (d), and aerobactin extracted ion count for the wild type and *schI* mutant (e).

After confirming the phenotypes caused by the knock-out of SchF0, SchE, SchH, or SchI, we quantified the relative abundances of each type of siderophore for each mutant (Figure 4) in order to establish the potential contribution of each siderophore to the siderome, as well as its potential contribution to iron chelation, in this organism. In the wild-type strain, serratiochelin represented over 80% of the siderophores and enterobactin, nearly 13%. There were small amounts of photobactin and aerobactin. In fact, the abundance of aerobactin was too low for quantification with the equipment used (Agilent single quadrupole mass spectrometer G6120a). Interestingly, knocking out SchI led to decreased production of serratiochelin (p=0.004) and photobactin (p=0.019). When the production of all polyamine siderophores was abolished (ΔSchH), the relative levels of enterobactin increased by ca. 50% (p = 0.004). The yields of all polyamine siderophores decreased when SchI was knocked out (p< 0.05), except for enterobactin, whose levels did not change. This suggests that the role of aerobactin in iron chelation in this organism is secondary when other siderophores are available. Moreover, it suggests that enterobactin takes up the iron chelation needs arising from lack of aerobactin. This compensatory activity by enterobactin may result from enterobactin requiring fewer enzymes and polyamines for assembly, in comparison with the polyamine siderophores.

**Figure 4.**
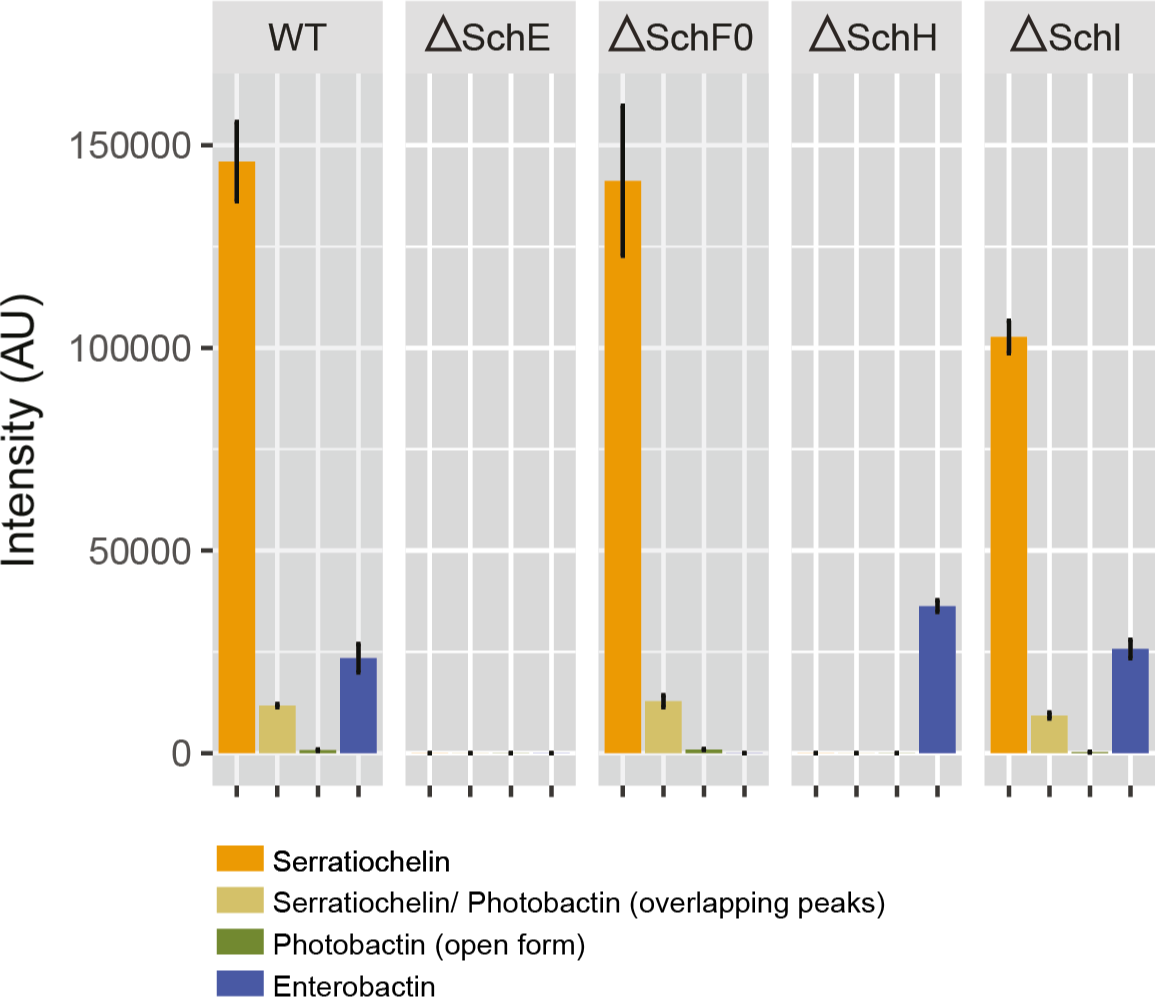
Relative abundance of serratiochelin, photobactin, and enterobactin in each mutant and wild-type strain. The low abundance of aerobactin did not allow for its relative quantification.

### Growth kinetics of *S. plymuthica* defective in the production of siderophores

Given that *S. plymuthica* synthetizes a plethora of siderophores, we were interested in understanding how each of these siderophores influences bacterial growth in iron-limited conditions. Therefore, we created mutant strains defective in the production of specific types of siderophores. Then we compared the growth of the mutants versus wild type, in the presence or absence of bipyridyl, a soluble iron chelator. Bipyridyl chelates any soluble iron that might still be present in the minimal medium and thus leads to the activation of the siderophore-producing machinery due to low soluble iron stress. We followed the growth of the strains over time and measured their maximum growth rate and maximum OD reached (Figure 5), in order to understand the relative importance of each group of siderophores (polyamine, catecholate, and hydroxamate siderophores). Overall, we found that the growth rate in minimal medium with bipyridyl was lower than in its absence for all strains except for the *schF0* mutant (Table 1). In the case of *schF0*, the maximum growth rate was, in fact, 39% (p=4.38 x 10^−7^) higher in the presence of bipyridyl than in its absence, but the maximum OD_610nm_ reached was 18% lower in the presence of bipyridyl than in its absence. We found that not producing catecholate or polyamine siderophores (SchE or SchH knockouts) increased the maximum growth rate of those mutants even in the presence of bipyridyl (Table 1). More precisely, there was an increase in growth rate of 30% for the SchH mutant and ca. 18% for the SchE mutant (p_SchH_=1.33 x 10^−35^, p_SchE_=4.43 x 10^−33^). Interestingly, when the organism was only incapable of producing enterobactin (SchF0 knockout), its maximum growth rate was reduced to 56% (without bipyridyl, p=3.90 x 10^−21^) and 89% (with bipyridyl, p=0.004) compared to wild-type. The maximum OD_610nm_ this mutant reached was also the lowest of all mutants and wild-type, even when grown in the absence of bipyridyl. To check whether these growth defects could be due to unexpected polar effects caused by the introduction of a suicide vector into *schF0*, we built a complementation strain, as well as related controls (Table 1). This complementation strain corresponds to the enterobactin-deficient strain (ΔSchF0) carrying a plasmid from which SchF0 is expressed. The complementation led the mutant strain to achieve both growth rates and ODs higher than the control (wild type carrying an empty pTrc99A plasmid) in the presence and absence of bipyridyl (p_GR_ = 3.51 x 10^−7^, p_OD_ = 3.96 ⨯ 10^−4^, and p_GR_ = 6.29 ⨯ 10^−14^ and p_OD_ = 1.96 ⨯ 10^−12^, respectively). The higher growth rates and ODs could be due to the plasmid being present in multicopy (pBR322 ori); if this was the case, SchF0 would have been more abundant in the complementation mutant than in the wildtype. These observations indicate that the slower growth of the SchF0 knockout strain can be attributed to its inability to produce enterobactin.

**Figure 5.**
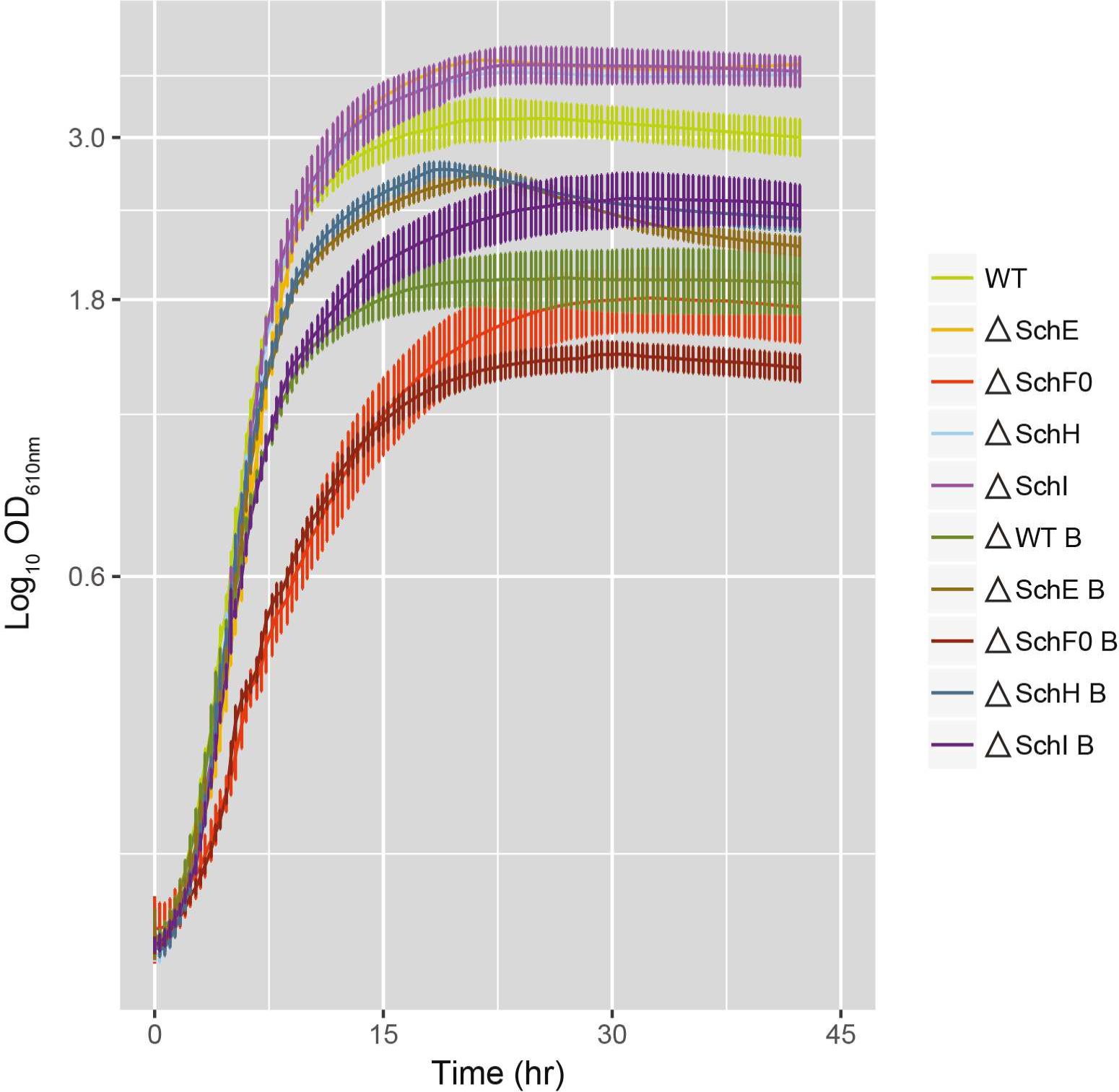
Time course of the OD_610nm_ for the mutant and wild-type strains, in the presence and absence of 0.1% bipyridyl (index B in the legend).

Our results suggest that although *S. plymuthica* produces more serratiochelin than any other siderophore, enterobactin and aerobactin seem to be the most cost-efficient siderophores, as revealed by the maximum OD_610nm_ and growth rate values for the SchE-, SchF0-, and SchI-deficient mutants (Table 1). In fact, enterobactin seems to play the most preponderant role in stimulating bacterial growth, as the lowest growth rates and OD_610nm_ were observed in the enterobactin-deficient strain.

### Characterization of the polyamine production superpathway

The biosynthesis of the *Serratia* spp.-proprietary siderophore serratiochelin is interconnected with that of polyamines. More precisely, the biosynthetic amide synthase SchH utilizes DAP as substrate for the assembly of serratiochelin (18). We also found that this same enzyme catalyzes the condensation of putrescine (rather than DAP) with DHB, to assemble photobactin (21) (Figure1). The natural biosynthesis of DAP is not a generalized feature of the Enterobacteriaceae, although its heterologous expression has been achieved in *E. coli* (25, 36). DAP is utilized in industry for the production of certain plastics (37–39) and as the basis for the production of agrochemicals (40). Homology searches for enzymes involved in the production of polyamines in *S. plymuthica* (41) enabled us to establish a putative amine production superpathway (Figure 6, Supplemental Table 2). We found that this organism encoded the machinery required for the synthesis of DAP, putrescine, cadaverine, and spermidine. Spermidine could potentially be synthesized from putrescine via S-adenosylmethionine decarboxylase, which has been found to transfer the aminopropyl group from S-adenosyl-3-(methylthio)propylamine to putrescine, originating spermidine in some prokaryotes (42–44) and eukaryotes (26, 45, 46). In some cases, spermidine can be converted to spermine by a second step that involves the transfer of an additional aminopropyl group to spermidine (47). Given that DAP and putrescine, although not present in the growth medium, are incorporated into serratiochelin and photobactin produced by *S. plymuthica*, it can be assumed that these polyamines are synthesized endogenously.

**Table 2.**
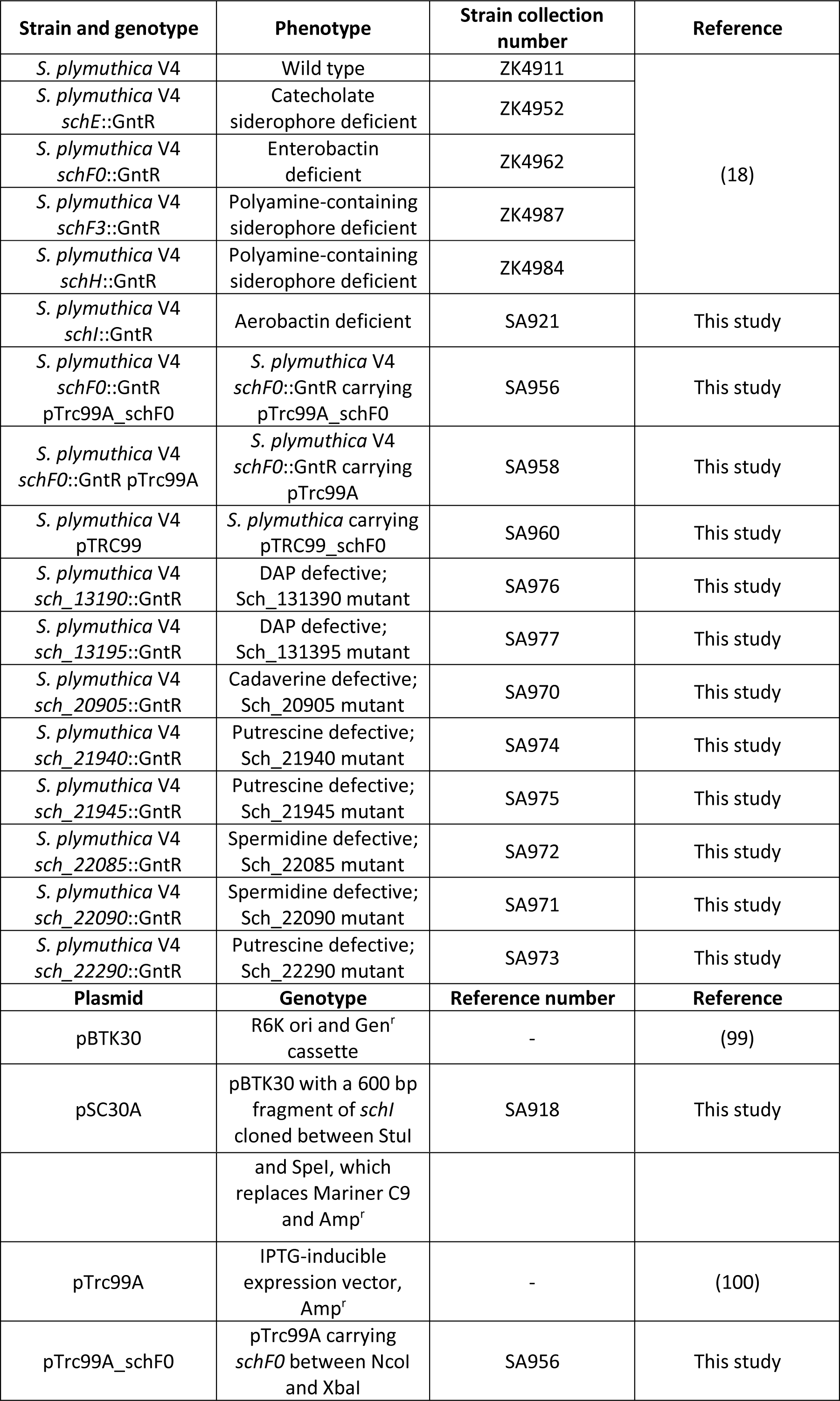
Strains and plasmids used in this study.

**Figure 6.**
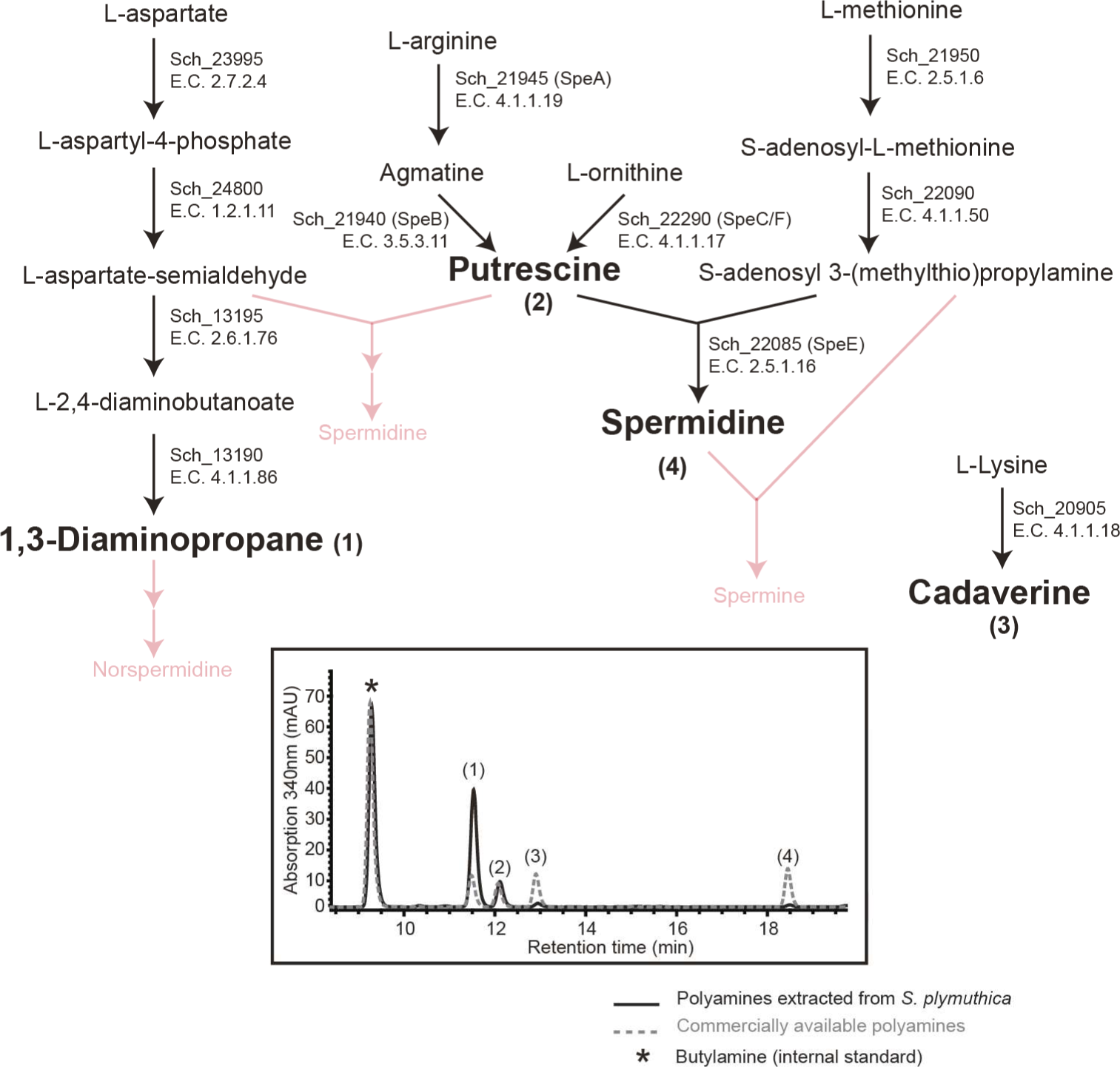
Proposed superpathway for polyamine production in *S. plymuthica* (top) and HPLC trace for the *S. plymuthica* samples and authentic standards (bottom). Polyamines not produced are displayed on the superpathway in red.

To determine which other polyamines were produced as well, samples obtained by the lysis of cell pellets were derivatized by dansylation and analyzed by tandem mass spectrometry for the presence of DAP, cadaverine, putrescine, spermidine, spermine, N-hydroxycadaverine, and aminopropylcadaverine. Furthermore, the abundance of DAP, cadaverine, putrescine, and spermidine was assessed by UV absorbance at λ=340nm in liquid chromatography based on polyamine standards purchased from Sigma - Aldrich (Supplemental Figures 5-8). *S. plymuthica* was found to produce 97.75±0.01 μmol/g DW of DAP and 30.83±0.003 μmol/g DW of putrescine, and small amounts of cadaverine (6.58±0.01 μmol/g DW) and spermidine (2.32±0.004 μmol/g DW). This level of DAP is ca. 60-fold higher than the highest reported yields of DAP naturally produced by other Proteobacteria (48).

Furthermore, part of the proposed polyamine production superpathway was confirmed by generating knockout strains and assessing their capacity to produce the predicted polyamines. The Sch_20905 knockout strain did not synthesize cadaverine (Supplemental Figure 9). We were unable to generate the other polyamine mutants, despite using the same approach as that used to generate all the other mutants, i.e., suicide vectors (see Methods). This suggests that DAP, putrescine, and spermidine may play essential roles in this organism. Spermidine, for example, is essential to the agrobactin-producing species *Agrobacterium tumefaciens* (49).

### Diversity of TonB-dependent receptors in *S. plymuthica*

Having elucidated the siderome of *S. plymuthica*, we were interested in understanding whether there was a corresponding TonB-dependent receptor (TBDR) for each type of siderophore produced. TBDRs are outer membrane proteins that, together with their inner membrane counterparts TonB, ExbB, and ExbD, transport selected siderophore-iron complexes, vitamin B12, nickel complexes, and carbohydrates into the cell (8). To assess the diversity of TBDRs, we queried the genome of *S. plymuthica* for known genes. As expected, we found the genes encoding the putative TBDRs specific for the siderophores produced, as well as others, for a total of 12 TBDRs. Specifically, we identified one putative receptor homologous to VuuA (vulnibactin) (50), ViuA (vibriobactin) (50), and PhuA (photobactin) (21), suggesting that a single receptor is capable of transporting all polyamine siderophores produced by *S. plymuthica*. Additionally, *S. plymuthica* encodes a FepA homolog that transports enterobactin (and colicins) (51), a LutA homolog that transports aerobactin (52), two CirA homologs that transport catecholate and colicin (53, 54), a YiuR homolog (55), and the homologous IrgA, which is a virulence factor without known transport functions (56). *S. plymuthica* was also found to encode receptors for fungal siderophores: a FhuE/PupB homolog that transports coprogen and rhodotorulic acid (57, 58), and a FhuA homolog that transports ferrichrome (59). In addition, it encodes receptors for mammalian hemoglobin, transferrin and lactoferrin (hemlactrns receptor family); hemin (HemR/HmuR/HxuC receptors) (60–62); and vitamin B12 and cobalamin (BtuB receptor) (63); as well as a homolog of the receptor BfrD/Fiu, which recognizes alcaligin, enterobactin, ferrichrome, and desferrioxamine B (64) (Figure 1, Supplemental Table 3). No serratiochelin-dedicated TBDR was found; thus, serratiochelin may be transported into the cell by the same TBDR as all other polyamine-containing siderophores (termed VuuA, ViuA, and PhuA, depending on the organism), as these siderophores are structurally related. Our analyses confirmed that, similar to other species, the genome of *S. plymuthica* encodes TBDRs that are more diverse than the siderophores this species synthesizes. This disparity is potentially associated with siderophore piracy, by which this species obtains iron via siderophores the organism did not spend energy making (15).

### Distribution of amide synthases across bacterial orders

Having established that *S. plymuthica* produces a diversity of polyamine-containing siderophores, we were interested in determining how widespread the distribution of the amide synthases is, as this could correlate with the discovery of new polyamine-containing siderophores. Amide synthases are enzymes crucial for the assembly of siderophores that contain polyamines (18, 28). These enzymes condense amines with other molecules, forming a carbon-nitrogen bond. To the best of our knowledge, the first amide synthase described as being involved in nonribosomal peptide assembly was VibH (28). VibH condenses norspermidine with DHB and is involved in the assembly of vibriobactin (28). SchH, a VibH homolog, is involved in the assembly of serratiochelin via DAP (18). PhbG, an uncharacterized homolog of VibH and SchH, is likely the amide synthase involved in the assembly of photobactin via putrescine in *Photorhabdus asymbiotica*, though this has yet to be experimentally confirmed. We then asked how widespread the distribution of amide synthases is and whether siderophores containing polyamines have already been characterized for those same organisms.

A tree containing 250 SchH homologs was generated using the Distance Tree of Results tool in BLASTp (Figure 7). Branches containing strains from the same species were collapsed for an easier interpretation of results. We then performed bibliographic searches aiming to find whether polyamine-containing siderophores had been characterized in these organisms. Of the organisms included in the tree, some have been described as producing nigribactin (*Vibrio nigripulchritudo*) (65), fluvibactin (*Vibrio fluvialis*) (66), vibriobactin (*V. cholerae*) (19), photobactin [*Photorhabdus* spp. (21) and *S. plymuthica* V4 (this study)], serratiochelin (*Serratia* spp.) (18, 67), parabactin (*Paracoccus* spp.) (68), and agrobactin (*Agrobacterium* spp.) (23). We anticipate that many more potentially new polyamine catechol siderophores have yet to be characterized for the remaining organisms, as they encode amide synthases as well as the remaining biosynthetic machinery for the assembly of catecholate siderophores. In the particular case of *Brucella* spp., brucebactin (69), its catechol siderophore, is unstable; this instability has prevented the elucidation of its structure.

**Figure 7.**
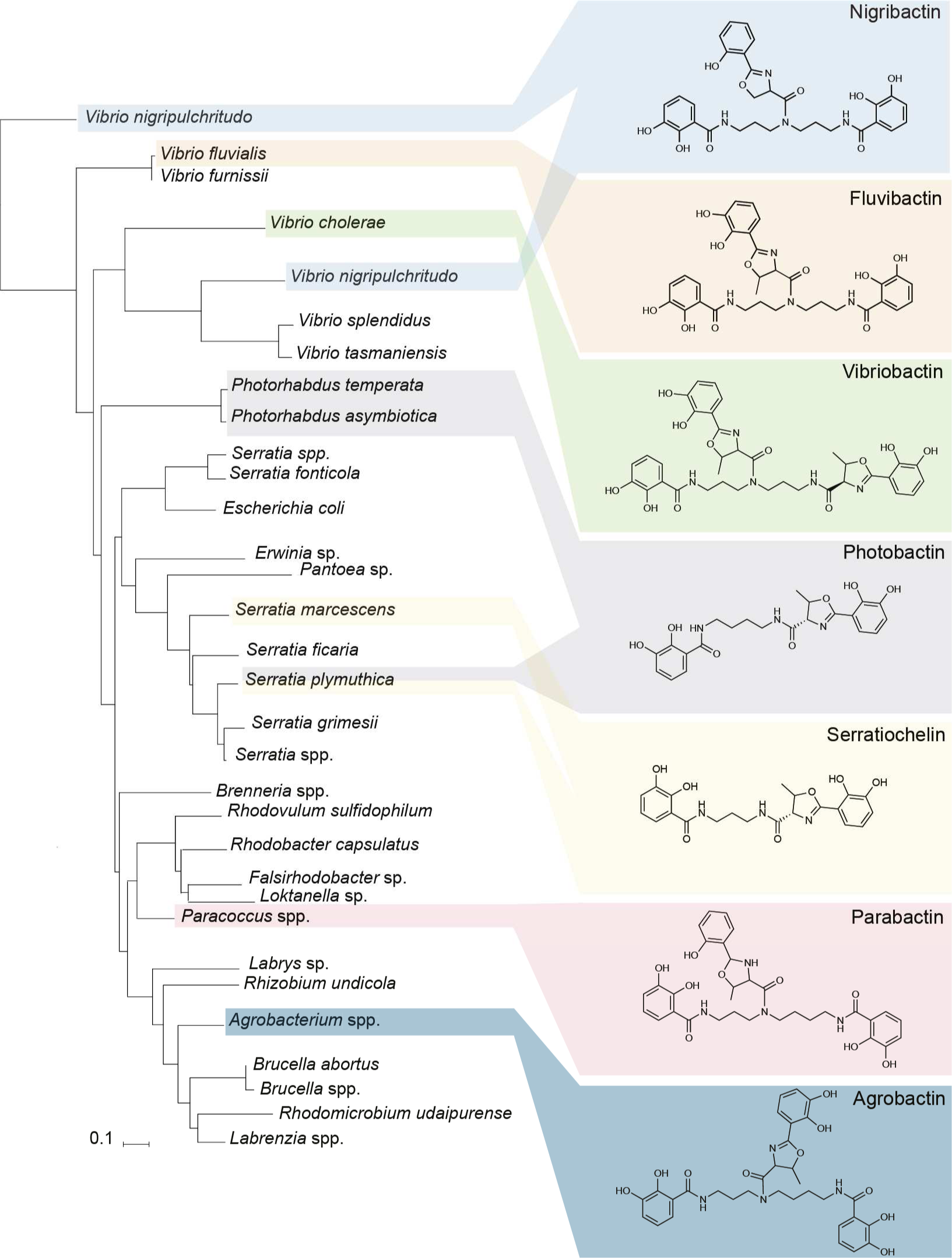
Phylogenetic tree for the distribution of amide synthases homologous to SchH across bacteria and their respective, known siderophores.

The analysis of the phylogenetic tree for SchH and its homologs did not reveal a particular evolutionary separation of the different molecules or of the aspects that make them different, such as the polyamine incorporation and condensation of DHBs on one or both primary amines (Figure 7; Supplemental Table 4). Amide synthases, key elements in the diversification of polyamine-containing siderophores, are encoded in diverse bacteria (Figure 7). Looking more closely at the standalone condensation domain of these amide synthases, we sought to correlate the active site residues (28), as well as the putative donor (polyamine) and acceptor binding (carrier protein-bound DHB) residues (70), with the polyamines they condensed. For all sequences, independently of the organism and polyamine-siderophore assembled, we found that the third residue in the active site motif HH**I**XXDG was conserved, whereas it usually is not conserved in condensation domains (HHXXXDG, Supplemental Table 4). For the donor (polyamine)-binding site, no clear sequence correlation was found. However, on the acceptor site, where DHB is presented by a carrier protein in all of the pathways, we found that the residue predicted to be crucial for substrate recognition (70) is invariably occupied by a valine (SchH residue 301). Furthermore, we identified a surface-exposed motif that appears to be characteristic of amide synthases and cannot be found in the condensation domains of other Nonribosomal Peptide Synthetases (71). As this motif is adjacent to W264 in VibH, a residue suggested to be important for protein-protein interaction (27), we propose that the entire motif supports the interaction with the carrier protein presenting DHB for condensation with the polyamine. To illustrate our findings, we generated a homology model for SchH based on the VibH crystal structure and mapped the described residues and motifs onto this model (Supplemental Figure 10).

Overall, our findings suggest that, similarly to what we observed with SchH, a single amide synthase might be capable of condensing a diversity of polyamines with the seemingly universal DHB acceptor. Nonetheless, this reaction might not be equally efficient for all polyamines, as we observed for putrescine and DAP, with the former being minimally used for siderophore assembly.

## Discussion

The capacity of pathogenic bacteria to acquire iron is intrinsically associated with their capacity to cause disease in humans (72). Siderophores, small molecules that act as iron scavengers and transporters, have thus been repeatedly categorized as virulence factors in some of the deadliest pathogenic bacteria that humans have encountered. Both *Yersinia pestis* and *Klebsiella pneumoniae*, for example, require the siderophore yersiniabactin for colonization of a mammalian host (73). It has been found that this same siderophore protects uropathogenic *E. coli* from copper toxicity during infection, while enterobactin allows the pathogen to survive (74). In fact, vertebrates do not have free, soluble iron (Fe^2+^) that bacteria can use for growth. This mechanism, termed nutritional immunity, is considered a vertebrate barrier to pathogen infection (75). Likely as a result of the pathogen-host struggle for iron, pathogens are thought to have evolved numerous ways of surviving in the host by resorting to siderophores to obtain iron. One of these strategies consists of expressing not one but a diversity of siderophores, as in the case of *S. plymuthica*. Although it rarely infects humans (76–78), *S. plymuthica* has collected several ways of synthesizing a diversity of siderophores and of acquiring xenosiderophores as well.

In this study, we found that *S. plymuthica* diversifies its siderophore production via a natural enzymatic mixing-and-matching. Initially this organism was characterized, like other *Serratia* species, as producing the siderophore serratiochelin (18, 67). In this study, we detected the biosynthesis of two additional catecholate non-ribosomal siderophores — enterobactin and photobactin — as well as aerobactin, a hydroxamate siderophore. The presence of these additional siderophores further supports the concept of the inheritance of genes as collectives (79); in other words, the concept that certain sets of genes evolve together and more quickly than their individual genes. The siderophores in *S. plymuthica* also exemplify how the enzymes encoded in these gene collectives can be diverted towards the assembly of not one but multiple molecules. Interestingly, this large diversity of siderophores in a single organism is more commonly associated with pathogenic bacteria than with environmental species such as *S. plymuthica*, which is rarely a cause of human disease (76). Apparently, the production of multiple siderophores is worth the metabolic cost for the bacteria that produce them, given the benefit of survival (80).

In fact, one of the most striking features of this organism is how efficiently it juggles two biosynthetic gene clusters to diversify the production of its catecholate siderophores. It was previously shown that the condensation domain-containing SchF0 is not involved in the assembly of serratiochelin (18). Indeed, SchF0 is not involved in the assembly of polyamine-containing siderophores, whereas it is crucial for the assembly of enterobactin in this organism. In *S. plymuthica*, instead of SchF0, three enzymes (SchF1F2F3) assemble serratiochelin and photobactin, while the enterobactin pathway provides DHB for these siderophores. These enzymes condense the thiol-bound DHB of the aryl carrier protein SchB with the thiol-bound threonyl of SchF3, instead of the seryl of SchF0, which is used for enterobactin. Thus, the diversity of the secondary metabolome may be underestimated, in particular if prediction tools are used, and too much reliance is placed on gene clustering for functional interpretation (81).

Another level of molecular diversification occurs when SchH condenses either DAP or putrescine with the acylated dihydroxybenzoyl of SchB, and SchF3 finalizes the assembly of serratiochelin (DAP) or photobactin (putrescine). *S. plymuthica* synthesizes serratiochelin and photobactin, which implies that it can synthesize the diamines DAP and putrescine. Queries to the genome of *S. plymuthica* for all the known genes that encode the enzymes involved in the polyamine superpathways of other organisms showed that the genetic makeup required was indeed present (Supplemental Table 2). The genetic basis of the superpathways was further confirmed by analyzing the polyamines actually produced by *S. plymuthica*, as assessed by LC-MS/MS detection and quantification of intracellular derivatized polyamine preparations (Figure 6).

The intercommunication of enzymes encoded by genes in two independent clusters, coupled with the substrate flexibility of SchH and the polyamine production profile of this organism, allows for two pathways to generate three distinct siderophores. To the best of our knowledge, the present study is the first to report that such interplay contributes to the diversification of nonribosomal peptides in a non-engineered organism. Furthermore, aerobactin, an additional, ribosomal siderophore, is also assembled independently of the others. Nonetheless, our results indicate that the production of polyamine siderophores might be particularly costly, as knocking out their production resulted in faster-growing cells, with cultures reaching higher ODs, as well (Table 1). Despite the abundance of siderophores produced, enterobactin still seems to play a major role in this organism: under iron-limiting conditions its absence leads to much slower growing cells that reach lower ODs. This suggests that the widespread presence of enterobactin among Proteobacteria might be due to its particularly high binding affinity for iron (K_d_=10^−52^ M)(82) and its cost efficiency.

The production of DAP in this organism, which is incorporated into its most abundant siderophore, was found to have the highest yield so far documented for a wild-type strain, to the best of our knowledge (25, 48, 83). This high yield suggests that bacteria that naturally produce polyamines and incorporate them into other molecules could potentially be optimized for the industrial-level production of polyamines. Polyamines are believed to be ancient molecules; not only are they present in all domains of life but there are multiple, convergent pathways resulting in a given polyamine (84, 85). However, it remains elusive how siderophores evolved and what selective forces gave rise to the intertwining of their biosynthetic pathways with polyamine biosynthesis. One hypothesis is that the competition for siderophores and the importance of preventing “cheaters” from utilizing xenosiderophores may be a driving force in siderophore evolution (11–13).

## Materials and Methods

### Strains, plasmids, and growth media

The strains and plasmids used and built for this study are listed in Table 2. Minimal medium optimized for the production of serratiochelins (18) was used for siderophore production. The minimal medium used consisted of Na2HPO4 (5.96 g/L), K2HPO4 (3.0 g/L), NH4Cl (1.0 g/L), NaCl (0.5 g/L), MgSO4 (0.058 g/L) and C6H12O6 (5.0 g/L). The final pH of the medium was 7.0.

### Siderome extraction and analysis

First, we grew *S. plymuthica* in the minimal medium described above. Overnight glucose-depleted cultures of *S. plymuthica* were used to inoculate 300 mL of minimal medium containing 0.1% bipyridyl. Bacteria were grown at 250 rpm with shaking at 30ºC, until glucose depletion (monitored using QuantoFix^®^ Glucose, Macherey-Nagel, USA). After the incubation period the cells were spun down and the supernatant was filter-sterilized (PES, 0.22 μm) and acidified with 0.1% trifluoracetic acid (TFA, final concentration). The acidified supernatant was run through Sep-Pak tC18 (200 mg) Reversed-Phase columns (Waters^®^), the columns were washed with 0.1% TFA in water, and the molecules were eluted with 95% acetonitrile acidified with 0.1% TFA as well. The extracted molecules, with varying hydrophobic character, were subsequently called “secondary metabolome,” in this article.

Liquid chromatography followed by tandem mass-spectrometry (LC-MS/MS) for secondary metabolome analysis was performed at the Small Molecule Mass Spectrometry core facilities at Harvard University. In order to detect and identify metabolites present in the samples, we performed XCMS Online analyses (86). The fragmentation patterns observed for the siderophores were compared with the described or predicted ones, utilizing ChemDraw.

### Generation of siderophore knockout mutants and determination of siderophore relative abundance

Based on prior knowledge of serratiochelin production, we asked whether these same gene clusters were responsible for the bioassembly of enterobactin and photobactin, detected in the siderome of *S. plymuthica* (18). We screened SchE (sch_19080), SchF0 (AHY08574.1), SchF3 (AHY05892.1), and SchH (AHY05888.1) knockout mutants for the production of each of these molecules. The mutants were kindly provided by Professor Roberto Kolter (Harvard Medical School).

The analysis of the metabolome also revealed the presence of aerobactin, a hydroxamate siderophore (34). In order to identify the operon responsible for the biosynthesis of aerobactin in *S. plymuthica*, we performed *iucA* homology searches in this organism using BLASTp (87, 88). *iucA* is one of the four biosynthetic genes in the aerobactin operon, previously characterized (34). This gene codes for a key enzyme in aerobactin biosynthesis, converting *N*^*6*^-acetyl-*N*^*6*^-hydroxy-L-lysine to *N*^1^-citryl-*N*^6^-acetyl-*N*^6^-hydroxy-L-lysine (34).

In order to search for i*ucA* in *S. plymuthica*, we generated a knockout mutant of *S. plymuthica* in which the gene *schI*, homologous to *iucA*, was disrupted by a suicide vector. This suicide vector was an R6K plasmid (which replicates only in the presence of the Pir protein) carrying a 350 bp region of homology towards the 5’ end of the gene. *schl* was disrupted to disable the biosynthesis of aerobactin by *S. plymuthica*. The suicide plasmid was cloned and maintained in *E. coli* S17-λPir-1, and the plasmid was moved to *S. plymuthica* by electroporation. The R6K origin does not replicate in *S. plymuthica* but integrates at a low rate into the chromosome at the designated locus of shared homology between the plasmid and the chromosome. The transformants were plated on selective medium, and the resulting colonies were PCR-verified for the integration of the suicide vector into *schI*.

A SchF0 knockout complementation mutant was also built, in order to confirm that the growth defects observed for the ΔSchF0 mutant resulted from the absence of SchF0 and not from polar effects. To create the complementation mutant, *schF0* was PCR-amplified from *S. plymuthica* V4 and cloned into plasmid pTRC99 by restriction digest and ligation. The construct was then electroporated into *S. plymuthica* V4 *schF0*::GntR. The empty vector pTRC99 was also electroporated into wild-type *S. plymuthica* V4, for use as control.

In order to characterize the sideromes of the wild-type strain and each of the mutants, we grew each strain in 300 mL of minimal medium supplemented with 0.1% bipyridyl, as described above. The cultures were monitored for glucose depletion and sampled at this point. Acidified (0.1% TFA) cell-free spent medium (270 mL) was supplemented with an internal control (0.5 mM Tyr-Tyr-Tyr eluent concentration, Sigma-Aldrich T2007). The spent medium was subsequently loaded into a Sep-Pak tC18 column (100 mg), as already described, and the compounds attached to the column were washed with 10% ACN (0.1% TFA) and eluted with 60%. Samples were then analyzed by High Performance Liquid Chromatography (HPLC) mass spectrometry (instrument: Agilent 1100, column: Agilent Zorbax Eclipse XDB-C18 80Å, 4.6 x 150 mm, 5μm; detector: Agilent single quadrupole mass spectrometer G6120a, injection volume 10 μL, gradient: 10% (v/v) ACN in water with 0.1% TFA for 1 minute, gradient to 55% ACN with 0.1% TFA over 25 minutes).

### Growth dynamics of the wild-type strain and siderophore mutants

Given the diversity of siderophores produced by *S. plymuthica*, we were interested in understanding the role they played in the survival of this strain in low iron conditions. We followed their growth kinetics, utilizing a microtiter plate reader programmed to take OD_610nm_ measurements every 20 minutes over 42 hours. Overnight (glucose depleted) cultures of the mutant and wild-type strains grown in minimal medium were used as inocula (OD_610nm_ 0.05).

Cultures of each strain (200 mL) were then incubated in 96-well plates, in the presence or absence of bipyridyl (6 wells per condition and strain, experiment repeated on 3 independent occasions). The data for each strain was averaged and plotted as OD_610nm_ as a function of time (hours). This data also enable the determination of the bacterial growth rate. The standard deviation for each group of data was calculated and is represented by error bars in the plots. The programming language R was used to calculate the maximum growth rate, the time at which maximum growth occurred (package *growthrates*), and the maximum OD_610nm_.

A separate experiment was performed to compare the growth kinetics of the *schF0* complementation strain *schF0*::GntR pTRC99_schF0 and wild-type pTrc99.

### Elucidation of the polyamine biosynthesis superpathway in *S. plymuthica* and polyamine production

*S. plymuthica* synthesizes serratiochelins by incorporating DAP into the nascent molecule. We thus asked what other polyamines this organism produces and whether they are utilized to generate analogs of serratiochelin.

We started by analyzing the superpathways for polyamine production in bacteria using MetaCyc (41). Using BLASTp (87), we queried *S. plymuthica* V4 for each of the enzymes in all three superpathways of polyamine biosynthesis. The similarity levels between the two homologous proteins were calculated using BLAST2p.

Aiming to confirm that the genes found indeed encoded enzymes involved in polyamine production, we tried to generate 8 knockout mutants using suicide vectors, as described for SchI. The genes we sought to disrupt were *sch_13190, sch_13195, sch_20905, sch_21940, sch_21945, sch_22085, sch_22090*, and *sch_22290*. Disruption of the genes *sch_21950, sch_23995*, and *sch_24800* was not attempted, as these genes have been deemed essential in *E. coli* (89). After multiple attempts and even after redesigning the suicide vector to increase regions of homology of >700 bp (in our hands, 400 bp suffice for single recombination in this strain), we were only able to disrupt *sch_20905*; therefore, we proceeded with this mutant alone.

To analyze cellular content for the polyamines predicted to be synthesized in the wildtype and mutant strains, cells were grown as described above, and upon glucose depletion 10 OD_610nm_ were pelleted. The pellets were resuspended in 500 μL of sterile water and 500 μL of 1.2 M perchloric acid (containing 1 mM butylamine as internal standard), vigorously vortexed in order to lyse the cells, and incubated for 1 hour at 37ºC. The lysate was centrifuged for 20 minutes at 4ºC and 12000 g, and the supernatant was collected (90).

Dansylation of polyamines was performed as described by Smith and Davies (91) with minor modifications, as follows: to 100 μL of the supernatant above, 200 μL of saturated NaCO_3_ (130 g/L) and 400 μL of dansyl-Cl (7.5 mg/mL acetone) were added. After incubation in the dark for 1 hour at 60ºC, 100 μL of proline (100 mg/mL) were added and the mixture was incubated for 30 minutes at 37ºC. Subsequently, the dansylated polyamines were extracted with 500 μL of toluene. For improved phase separation, the samples were centrifuged for 3 minutes at 3000 g. The organic phase was dried under a nitrogen stream and the pellet was resuspended in 50 μL of methanol. For identification of the polyamines produced, a 10 μL aliquot was injected into a high-resolution, accurate mass Q Exactive Plus Orbitrap, with positive ionization and mass scan ranging from 66 to 990 m/z (resolution 17.500 FWHM), and separated over the course of 30 minutes at a flow rate of 0.8 mL/min, with a gradient of 10% ACN in H_2_O to 100% ACN.

Authentic standards 1,3 - diaminopropane, putrescine, spermidine, spermine, and cadaverine were acquired from Sigma - Aldrich and used for determination of their fragmentation pattern, for comparison with the test samples. The standards were dansylated at the same time as the samples. The samples were analyzed as described elsewhere (91), at the Small Molecule Mass Spectrometry core facilities at Harvard University.

For quantification of the polyamines in *S. plymuthica*, samples were prepared as mentioned above except that they were resuspended in 100 μL of MeOH and analyzed by HPLC (instrument: Agilent 1100, column: Agilent Zorbax Eclipse XDB-C18 80Å, 4.6 x 150 mm, 5μm; detector: Agilent diode array detector G1315B, λ=340nm, injection volume 25 μL, gradient: 60% (v/v) MeOH in water for 1 minute, gradient to 100% MeOH over 23 minutes). Integrated peak areas were normalized based on the internal standard and converted to concentrations in mM, based on three samples of known concentration of each authentic standard.

The dry weight was determined as the pellet biomass after 24 hours at 85ºC (92), and used to calculate the concentration in mole per gram of dry weight.

### Comparative analysis of amide synthase

In order to determine the conservation and distribution of amide synthases, we queried the NCBI database for 250 homologs of SchH. In order to analyze how these homologs clustered together, based on protein sequence similarity, we downloaded the respective Newick trees (Neighbor Joining, 0.85 maximum sequence difference, Grishin distance).

The condensation domain of VibH has been thoroughly characterized by others, who revealed that its active site contains the highly conserved motif HHXXXDG (27, 93, 94). We asked whether the three variable residues correlated with the polyamine condensed into the nascent molecule. For this we queried the NCBI sequence database for homologs of the amide synthase SchH, from genera known to include strains that synthesize polyamine-containing siderophores. These were *S. plymuthica* (serratiochelin and photobactin), *Serratia marcescens* (serratiochelin), *Paracoccus* spp. (parabactin), *Agrobacterium*/*Rhizobacterium* (agrobactin), *Vibrio cholerae* (vibriobactin), *Vibrio fluvialis* (fluvibactin), *Vibrio nigripulchritudo* (nigribactin) and *Vibrio vulnificus* (vulnibactin). The available sequences (up to 250 per genus or strain) were aligned in NCBI (gap penalties −11, −1; end-gap penalties −5, −1; maximum cluster distance 0.8). The alignments were downloaded, further processed in CLC Sequence Viewer 7, and queried for the active site sequence. The residue variation in the conserved motif per bacterial species known to produce polyamine-containing siderophores was analyzed.

### Identification of TonB-dependent siderophore receptors in *S. plymuthica* V4

Having established that *S. plymuthica* V4 produced a large repertoire of siderophores, we then asked whether this organism has a corresponding diversity of TonB-dependent siderophore receptors. In order to determine the diversity of TBDRs encoded in the chromosome of *S. plymuthica* V4, we queried it for each of the TBDR families thus far characterized (Uniprot reviewed entries only). The TBDRs found were compared to their respective protein reference sequences using NCBI’s BLAST2p tool. This tool aligns two proteins and computes their level of similarity (88).

### Homology modeling

To generate a homology model for the amide synthase SchH, we used the SWISS-MODEL server (95–97) with the VibH crystal structure as input (PDB: 1l5a, chain A; quality assessment: QMEAN −3.2). The model was visualized using PyMol (98).

### Statistical treatment of data

Statistical significance of the results was analyzed using the unpaired, unequal variance non-parametric t-Test. Siderophore and polyamines relative and absolute levels, respectively, were determined from 3 independent experiments, with technical duplicates. The growth curves were calculated with the data obtained from 3 independent experiments, with 6 technical replicates. The statistical significance is represented in the figures by * (p<0.050), ** (p<0.010) or *** (p<0.001).

## Acknowledgments

This research was supported by the NIH Director’s New Innovator Award 1-DP2-OD008435-01 to T.K.L., Novartis Award 68148476 to T.K.L., the United States Defense Threat Reduction Agency (HDTRA1-14-1-0007 and HDTRA1-15-1-0050) to T.K.L., and the Office of Naval Research (ONR 4500000552) to T.K.L. K.H, is grateful for the support by the Human Frontier Science Program (Grant Number LT000969/2016-L).

