## Supporting Information for "Natural combinatorial genetics and prolific polyamine production enable siderophore diversification in Serratia plymuthica"

* Timothy K. Lu

Contents

Supplemental Tables……………………………………………………………………………2

Supplemental Figures…………………………………………………………………………6

#### Supplemental Tables

**Supplemental Table 1.** Enzymes involved in the biosynthesis of the hydroxamate siderophore aerobactin, identified by their protein name and locus in the chromosome of *S. plymuthica* V4, and compared to enzymes from other aerobactin-producing bacteria on amino acid level.

| Protein | Comparison | Positives | E-value | Identity | Gaps |
|---|---|---|---|---|---|
| IucD/SchL (sch_04435) | *Y. pekkanenii* WP_049612679.1 | 93% | 0.0 | 89% | 0% (n=0) |
| | *E. coli* KRT38817.1 | 80% | 0.0 | 70% | 0% (n=0) |
| | *E. aerogenes* CZY40742.1 | 81% | 0.0 | 71% | 0% (n=0) |
| IucC/SchK (sch_04440) | *Y. pekkanenii* WP_049612681.1 | 91% | 0.0 | 87% | 0% (n=0) |
| | *E. coli* KRT38816.1 | 84% | 0.0 | 72% | 0% (n=2) |
| | *E. aerogenes* SAC04707.1 | 83% | $5.0 \times 10^{-166}$ | 72% | 0% (n=2) |
| IucB/SchJ (sch_04445) | *Y. enterocolitica* WP_050143497.1 | 91% | 0.0 | 85% | 0% (n=0) |
| | *E. coli* KRT38815.1 | 80% | $5.0 \times 10^{-163}$ | 69% | 0% (n=0) |
| | *E. aerogenes* SAC04673.1 | 81% | $5.0 \times 10^{-166}$ | 70% | 0% (n=0) |
| IucA/SchI (sch_04450) | *Y. enterocolitica* WP_050143497.1 | 91% | 0.0 | 85% | 0% (n=0) |
| | *E. coli* KRT38814.1 | 82% | 0.0 | 73% | 0% (n=0) |
| | *E. aerogenes* SAC17751.1 | 83% | 0.0 | 73% | 0% (n=0) |

**Supplemental Table 2**. *S. plymuthica* proteins involved in the conversion of specific amino acids to polyamines and comparison to reference sequences on amino acid level.

| Enzyme activity | Protein | Reference sequence | Max score | Total score | E-value | Identity |
|---|---|---|---|---|---|---|
| Aspartate kinase | Sch_23995 | WP_015962105.1 | 796 | 796 | 0.0 | 89% |
| Aspartate-semialdehyde dehydrogenase | Sch_24800 | WP_009111079.1 | 696 | 696 | 0.0 | 91% |
| Diaminobutyrate-2-oxoglutarate aminotransferase | Sch_13195 | WP_017802287.1 | 894 | 894 | 0.0 | 94% |
| 2,4-diaminobutyrate decarboxylase | Sch_13190 | WP_014543183.1 | 933 | 933 | 0.0 | 92% |
| Arginine decarboxylase | Sch_21945 | WP_019082398.1 | 1240 | 1240 | 0.0 | 91% |
| S-adenosylmethionine synthetase | Sch_21950 | WP_005123306.1 | 764 | 764 | 0.0 | 95% |
| Agmatinase | Sch_21940 | WP_008499737.1 | 566 | 566 | 0.0 | 88% |
| Ornithine decarboxylase | Sch_22290 | WP_006327912.1 | 1500 | 1500 | 0.0 | 99% |
| Spermidine synthase | Sch_22085 | Q66EH3.1 | 552 | 552 | 0.0 | 91% |
| S-adenosylmethionine decarboxylase | Sch_22090 | Q66EH4.1 | 509 | 509 | 0.0 | 91% |
| Lysine decarboxylase | Sch_20905 | WP_013814260.1 | 1489 | 1489 | 0.0 | 99% |

**Supplemental Table 3.** Putative TonB-dependent receptor homologs found in the chromosome of *S. plymuthica*, molecule transported, and level of similarity with the reference sequence.

| TonB-dependent receptors | Putative homolog | Homolog reference sequence | Molecule transported | Max score | Total score | Identity |
|---|---|---|---|---|---|---|
| FhuE/PupB/FptA/FpvA/PbuA/PupA | Sch_19445 | WP_012146208.1 | Coprogen, rhodotorulic acid, pseudobactin | 1375 | 1375 | 91% |
| FhuA/FatA/Fct | Sch_21980 | WP_004947851.1 | Ferrichrome, ferrichrysobactin, photobactin | 1509 | 1509 | 99% |
| Fep/Cbt/ Cbr/FeuB/FepA/PfeA/IroN/BfeA | Sch_19130 | WP_006327062.1 | Enterobactin; colicins B, D | 1562 | 1562 | 99% |
| BfrD/Fiu/FoxA | Sch_14815 | WP_004706957.1 | Ferrioxamine B | 1057 | 1057 | 70% |
| Cir/IrgA/YiuR | Sch_18005 | WP_016926997.1 | Catecholate, colicin | 1129 | 1129 | 83% |
| CirA/YncD | Sch_15725 | WP_006321367.1 | Catecholate, colicin | 1447 | 1447 | 98% |
| | Sch_02800 | WP_006323436.1 | | 1533 | 1533 | 100% |
| VuuA/ViuA/PhuA | Sch_04725 | WP_017891746.1 | Vulnibactin, vibriobactin, photobactin, serratiochelin? | 1377 | 1377 | 94% |
| IutA/RhtA | Sch_04430 | WP_002211646.1 | Aerobactin/rhizobactin | 1327 | 1327 | 87% |
| Hemlactrns | Sch_12995 | WP_006318362.1 | Hemoglobin, transferrin, lactoferrin | 1575 | 1575 | 97% |
| HemR/HmuR/HxuC | Sch_11745 | WP_011815406.1 | Hemin | 1036 | 1036 | 70% |
| BtuB | Sch_25400 | | Vitamin B12/cobalamin | | | |

Maximum score = Total score

E-value 0.0 for all sequences

**Supplemental Table 4.** Amide synthase active site residues by species and polyamine condensed, based on NCBI sequence data and characterized siderophores.

| **Species (number of sequences)** | **Variable residues in active site motif (% of sequences)** | **Proposed acceptor binding motif (% of sequences)** | **Polyamine** |
|---|---|---|---|
| *S. plymuthica* (n=9) | III (100%) | MNRRSS (100%) | Diaminopropane/putrescine |
| *S. marcescens* (n=64) | ILV (86%) ILI (14%) | MNRRGQ (77%) LNRRGQ (20%) LNRRGH (3%) | Diaminopropane |
| *Photorhabdus* spp. (n=9) | ILL (89%) IIL (11%) | MNRRTR (100%) | Putrescine |
| *Agrobacterium/Rhizobacterium* (n=32) | IVV (97%) IIV (3%) | MNRRSS (100%) | Spermidine |
| *Paracoccus* spp. (n=9) | IAL (100%) | MSRMGS (88.9%) LSRMGS (11.1%) | |
| *Vibrio cholerae* (n=90) | IVL (98%) IIL (2%) | MNRWGS (100%) | Norspermidine |
| *Vibrio fluvialis* (n=13) | IVM (100%) | MSRWGS (100%) | |
| *Vibrio nigripulchritudo* (n=11) | IAV (64%) IVL (36%) | MNRFGS (37%) MNRMGN (63%) | |
| *Vibrio vulnificus* (n=42) | IAL (100%) | LNRMGN (100%) | |

### Supplemental Figures

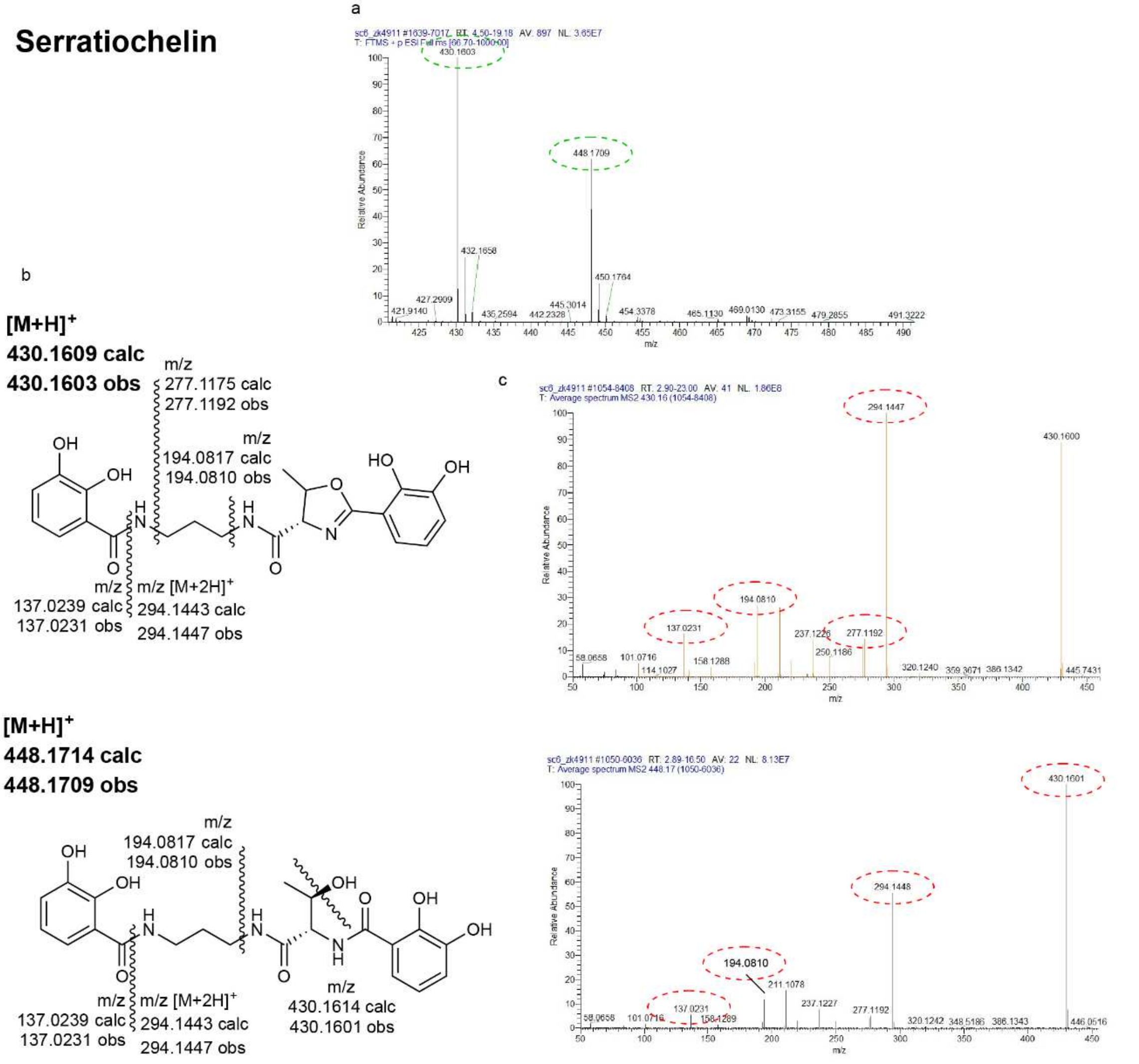


**Supplemental Figure 1.** ESI-MS for open and closed-ring serratiochelin (a), their structures, calculated and observed mass (b) and observed ESI-MS/MS (c). The fragment masses indicated (b) correspond to those observed, circled in red (c).

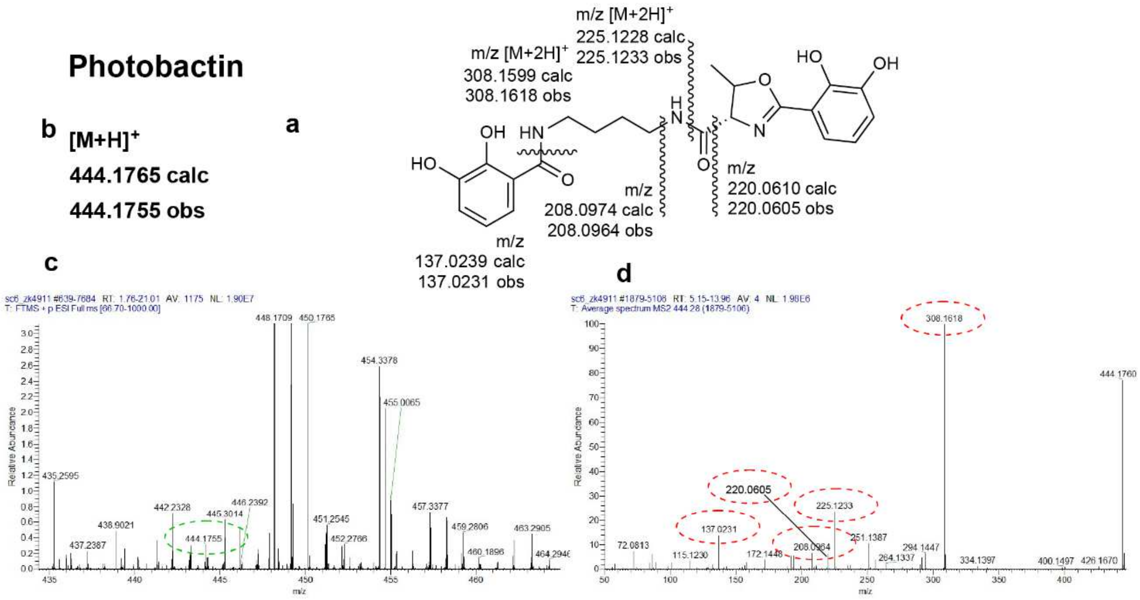


**Supplementary Figure 2**. Proposed structure and fragmentation pattern (**a**), calculated and observed mass (**b**), ESI-MS (**c**) and ESI-MS/MS (**d**) for M5Tc. In **c**, the mass value circled in green highlights the observed [M+H]+ that is consistent with the calculated one. In **d**, the mass values resulting from the fragmentation of the molecule circled in **c**, which are consistent with the expected pattern of fragmentation shown in **a**, are circled in red.

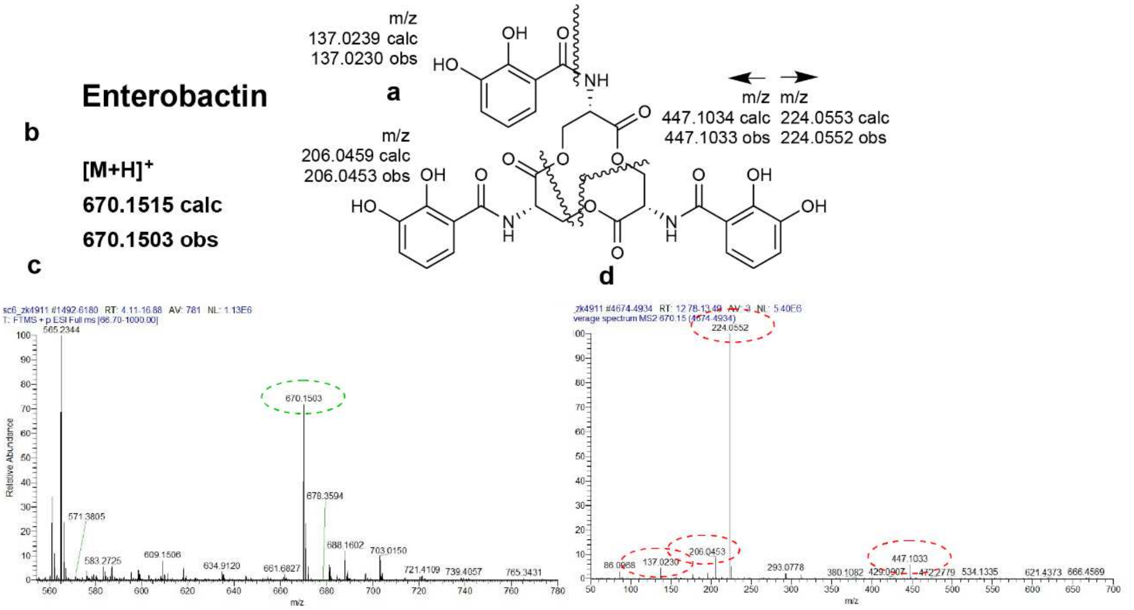


**Supplementary Figure 3**. Proposed structure and fragmentation pattern (**a**), calculated and observed mass (**b**), ESI-MS (**c**) and ESI-MS/MS (**d**) for enterobactin. In **c**, the mass value circled in green highlights the observed [M+H]+ that is consistent with the calculated one. In **d**, the mass values resulting from the fragmentation of the molecule circled in **c**, which are consistent with the expected pattern of fragmentation shown in **a**, are circled in red.

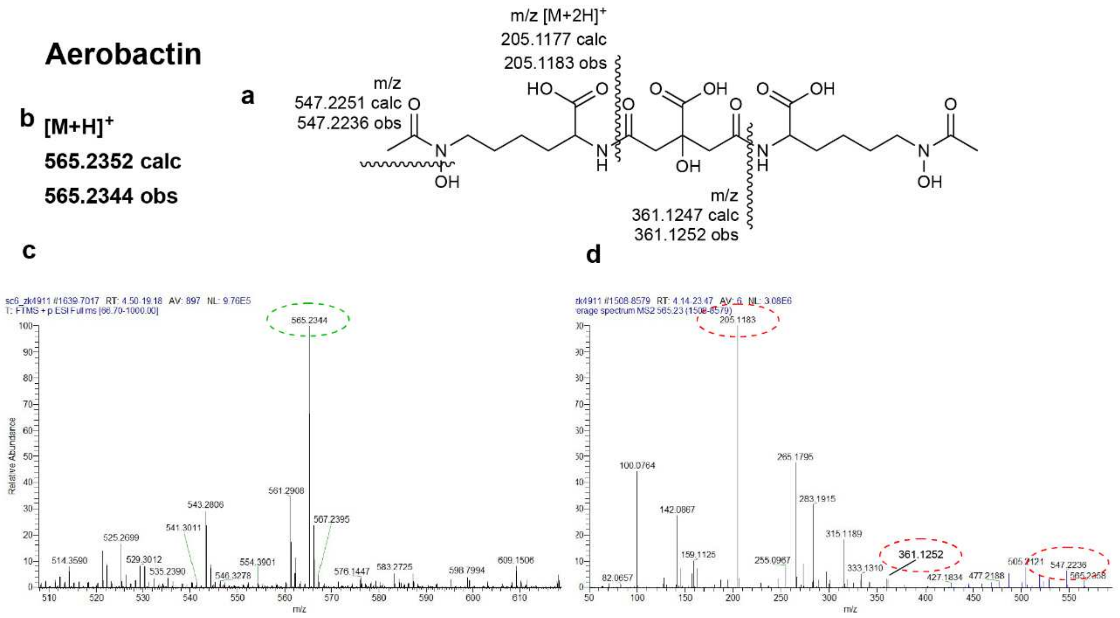


**Supplementary Figure 4**. Proposed structure and fragmentation pattern (**a**), calculated and observed mass (**b**), ESI-MS (**c**) and ESI-MS/MS (**d**) for aerobactin. In **c**, the mass value circled in green highlights the observed [M+H]+ that is consistent with the calculated one. In **d**, the mass values resulting from the fragmentation of the molecule circled in **c**, which are consistent with the expected pattern of fragmentation shown in **a**, are circled in red.

(a)

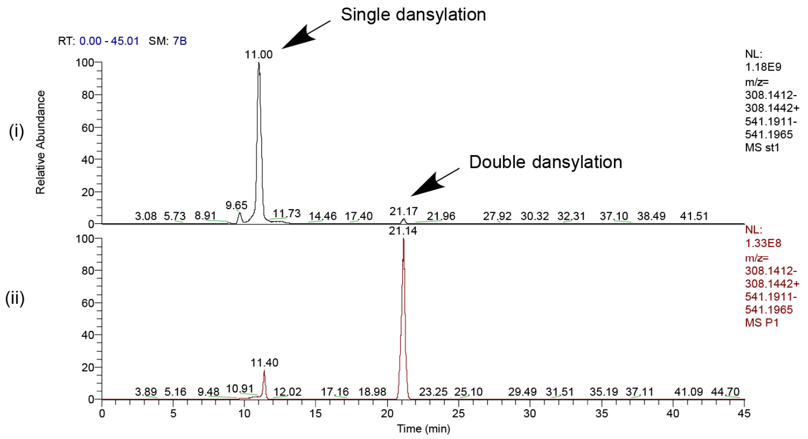


(b)

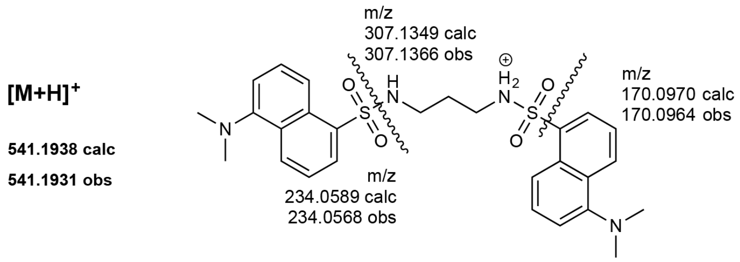


(c)

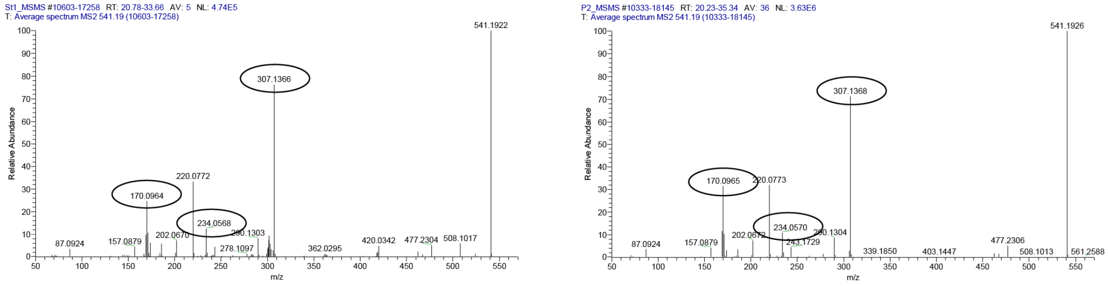


(i) (ii)

**Supplemental Figure 5**. Dansylated 1, 3 - Diaminopropane: (a) extracted ion count for partially or fully dansylated polyamines in the control (i) and test sample (ii); (b) structure, exact mass and fragmentation pattern for the standard (i) and test sample (ii).

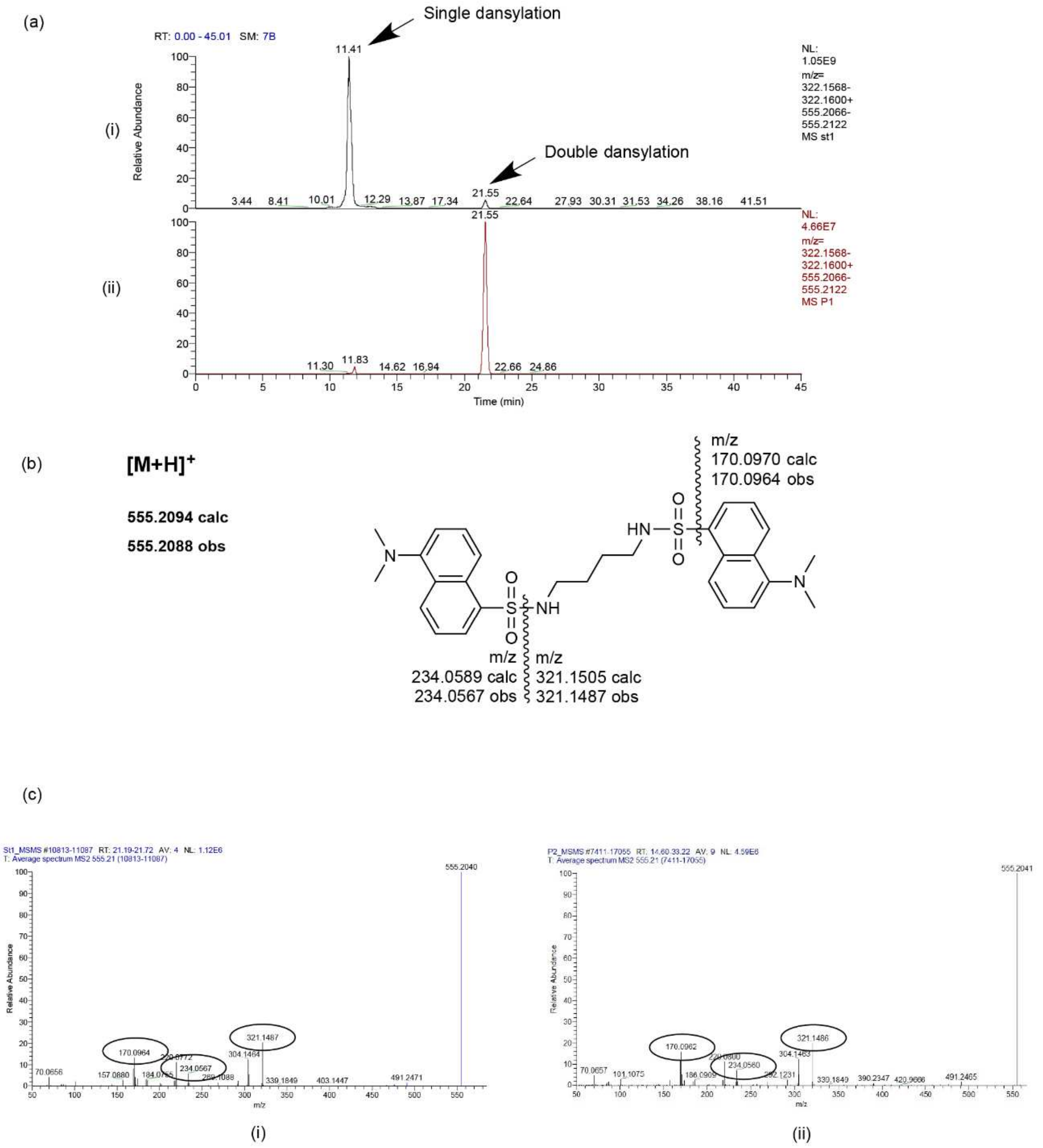


**Supplemental Figure 6**.:Dansylated putrescine: (a) extracted ion count for partially or fully dansylated polyamines in the control (i) and test sample (ii); (b) structure, exact mass and fragmentation pattern for the standard (i) and test sample (ii).

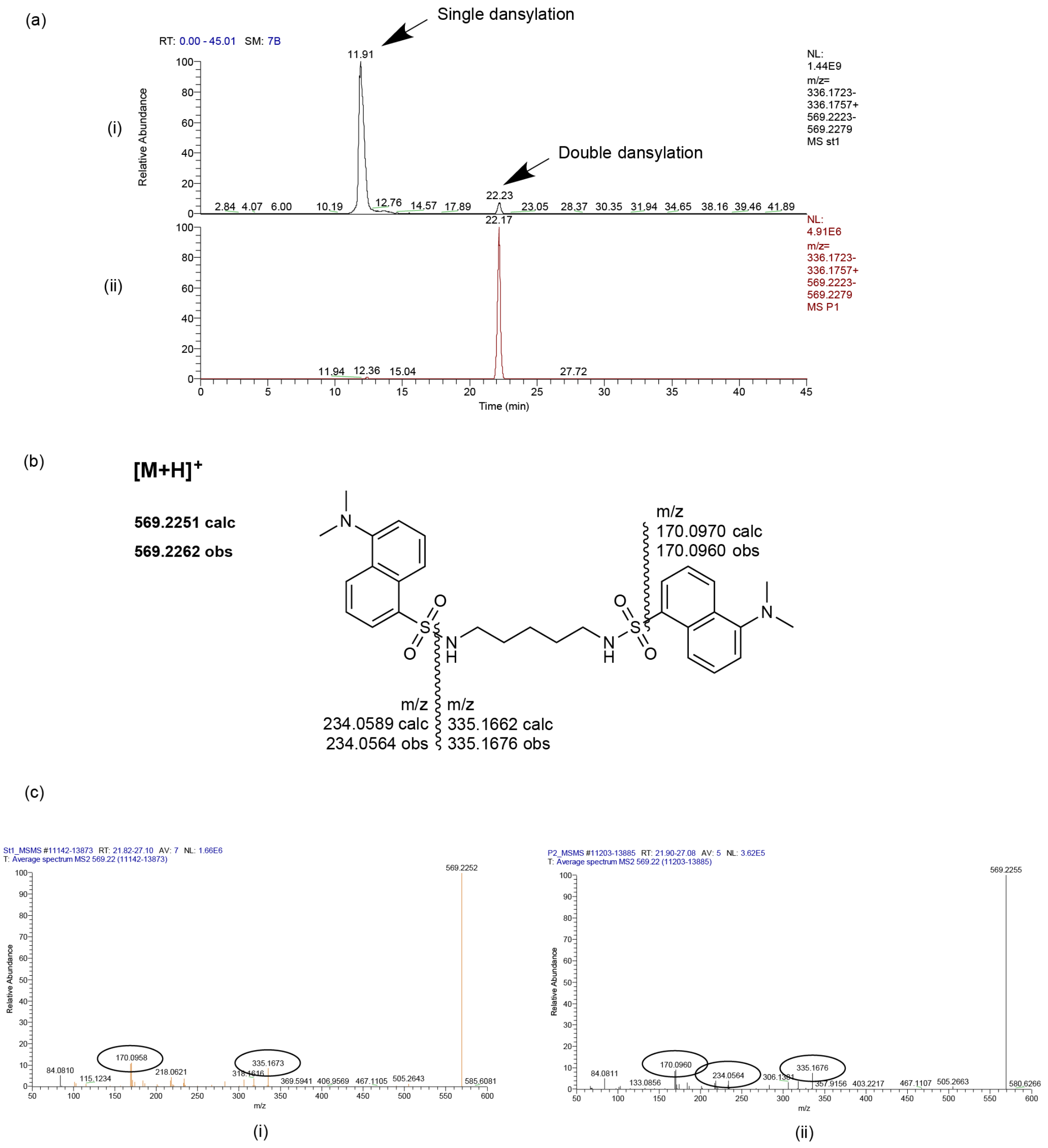


**Supplemental Figure 7.** Dansylated cadaverine: (a) extracted ion count for partially or fully dansylated polyamines in the control (i) and test sample (ii); (b) structure, exact mass and fragmentation pattern for the standard (i) and test sample (ii).

(a)

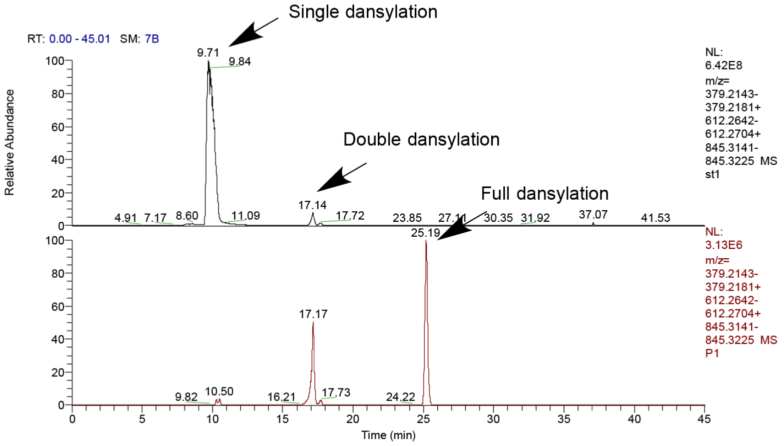


(b)

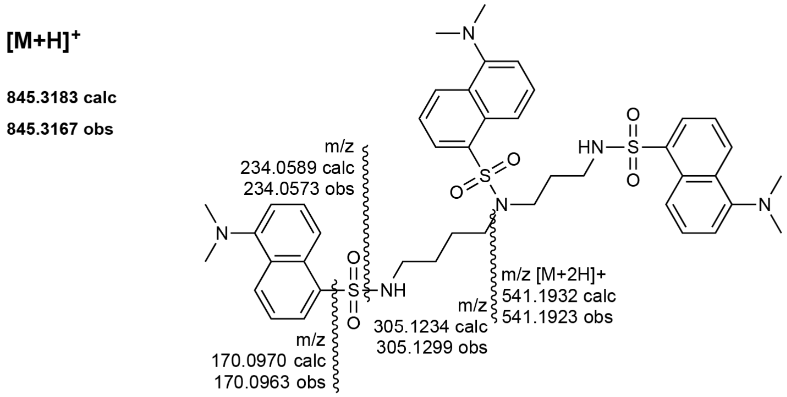


(c)

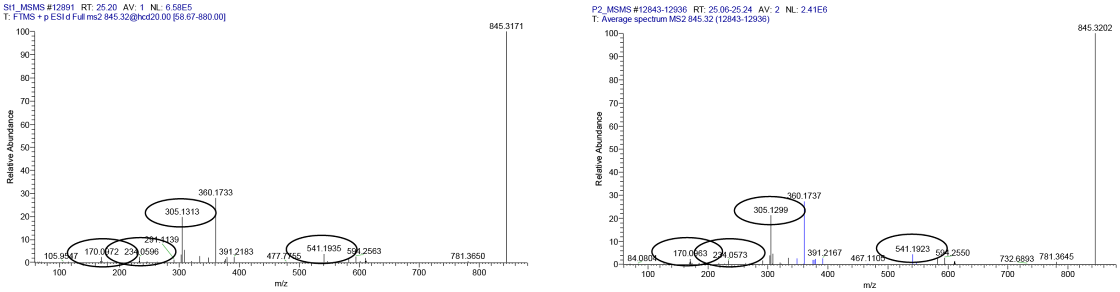


(i) (ii)

**Supplemental Figure 8.** Dansylated spermidine: (a) extracted ion count for partially or fully dansylated polyamines in the control (i) and test sample (ii); (b) structure, exact mass and fragmentation pattern for the standard (i) and test sample (ii).
